# Biliverdin reductase bridges focal adhesion kinase to Src to modulate synaptic signaling

**DOI:** 10.1101/2020.12.01.405720

**Authors:** Chirag Vasavda, Evan R. Semenza, Jason Liew, Ruchita Kothari, Ryan S. Dhindsa, Shruthi Shanmukha, Cristina Ricco, Robert Tokhunts, Adele M. Snowman, Lauren Albacarys, Bindu D. Paul, Solomon H. Snyder

## Abstract

Synapses are complex bridges that connect discrete neurons into vast networks that send, receive, and encode diverse forms of information. However, they must remain dynamic in order to adapt to changing inputs. Here, we report that the enzyme biliverdin reductase (BVR) physically links together key molecules in focal adhesion signaling at the synapse. In challenging mice with a battery of neurocognitive tasks, we first discover that BVR null (BVR^-/-^) mice exhibit profound deficits in learning and memory. We uncover that these deficits may be explained by a loss of focal adhesion signaling that is both transcriptionally and biochemically disrupted in BVR^-/-^ hippocampi. We learn that BVR mediates focal adhesion signaling by physically bridging the initiatory kinases FAK/Pyk2 to the effector kinase Src. Activated Src normally promotes synaptic plasticity by phosphorylating the N-methyl-D-aspartate (NMDA) receptor, but FAK/Pyk2 are unable to bind and stimulate Src without BVR. Src itself is a molecular hub upon which many signaling pathways converge in order to stimulate NMDA neurotransmission, positioning BVR at a prominent intersection of synaptic signaling.

## Introduction

The molecules that lie within synapses are the substrates for some of the most essential functions of neurons. Receptors at synapses receive and transduce signals from adjacent neurons, channels influence the excitability of the pre- and post-synaptic neuron, and adhesion molecules make and break new connections. Together, these molecules build complex bridges that connect discrete neurons into vast networks that send, receive, and encode diverse forms of information.

Synapses are plastic structures that are dynamically regulated at multiple levels in order to adapt to new information. Synapses can strengthen or weaken over time, and such activity-dependent changes in synaptic signaling play an important role in the development and maintenance of neural circuits (*1*). Depending on the inputs into a synapse, the underlying actin cytoskeleton can expand or shrink (*2*), channels may be inserted or endocytosed (*3*), or new proteins can be translated (*4*). This plasticity is thought to drive learning and memory, encoding them by changing the vastly interconnected neural circuits in the brain (*5*). Long-term potentiation (LTP) is one phenomenon in which certain synaptic inputs, such as high-frequency stimulation, result in a persistent, enduring increase in the strength of synaptic transmission (*6*). It is thought that the molecular processes that underlie LTP are principal components of synaptic plasticity during learning (*7–9*).

LTP was first discovered in the hippocampus (*10*) but is now known to occur throughout the brain, including the cortex, amygdala, and cerebellum (*11*). The exact mechanics of LTP can vary among these regions and can manifest differentially depending on a number of factors, but LTP is classically mediated by the *N*-methyl-D-aspartate (NMDA) receptor (*6*). The NMDA receptor is a ligand-gated ion channel activated by the excitatory neurotransmitter glutamate. It is unique among ionotropic glutamate receptors in that it is permeable to calcium as well as sodium and potassium, whereas the others only gate sodium and potassium. Calcium influx through NMDA receptors is central to LTP and initiates a series of biochemical signals, including activation of calcium-calmodulin and protein kinase C (PKC) (*12*). NMDA receptors also mediate redox signaling by triggering synthesis of nitric oxide and superoxide (*13, 14*). Alongside calcium, nitric oxide and superoxide are key to inducing LTP in the hippocampus (*15, 16*).

Recently, we discovered that the heme metabolite bilirubin influences NMDA redox signaling by scavenging superoxide (*17*). Mice lacking the enzyme that synthesizes bilirubin, biliverdin reductase (BVR), exhibit dysregulated NMDA redox signaling. BVR null mice (BVR^-/-^) exhibit exaggerated locomotor activity when injected with NMDA or MK-801 (an NMDA receptor agonist and antagonist, respectively), further suggesting that BVR^-/-^ mice exhibit irregular NMDA receptor signaling. Here, we considered whether losing BVR impacts NMDA receptor-dependent synaptic signaling. By leveraging a combined behavioral, computational, and biochemical approach, we discover that BVR^-/-^ mice exhibit profound learning and memory deficits. We find that BVR physically interacts with key molecules in focal adhesion signaling to mediate synaptic signaling in the hippocampus.

## Results

### BVR^-/-^ mice exhibit impaired learning and memory, but no anxiety or despair

To determine whether eliminating BVR impacts learning and memory, we evaluated the performance of BVR^-/-^ mice performed against their wild-type littermates in several neurobehavioral assays. We first assessed whether BVR^-/-^ mice exhibit spatial learning deficits in the Morris water maze. The Morris water maze reliably correlates with hippocampal synaptic plasticity (*18*), and animals with lesions in the dentate gyrus or subiculum perform poorly (*19, 20*). In this assay, mice are trained over four days to locate a submerged platform in a pool of cloudy water. Each day, we begin by revealing the platform to the mice with unique visual cues prominently displayed on the walls of the pool. We then place the mice at different locations around the pool four times and monitor how long it takes them to re-locate the platform as a measure of their spatial learning. Mice with deficits in spatial learning require more time to find the platform throughout training. After four days of training, the mice are tasked once again with locating the platform but without being shown it beforehand. Mice with spatial learning and memory deficits will take more time to reach the platform on average than those that remember where to travel. In training, WT mice quickly learn by days 3 and 4 where the platform lies whereas BVR^-/-^ mice are unable to locate it (**Fig. 1a**). BVR^-/-^ mice also spend over twice as long on average locating the platform than WT mice after four days of training, with some BVR^-/-^ mice never finding the platform (**Fig. 1b**). BVR^-/-^ mice are not just slower to reach the platform because they are weaker swimmers, since BVR^-/-^ mice require more time to locate the platform despite swimming faster and covering a greater distance than WT mice (**Fig. 1c-d**).

**Figure 1.**
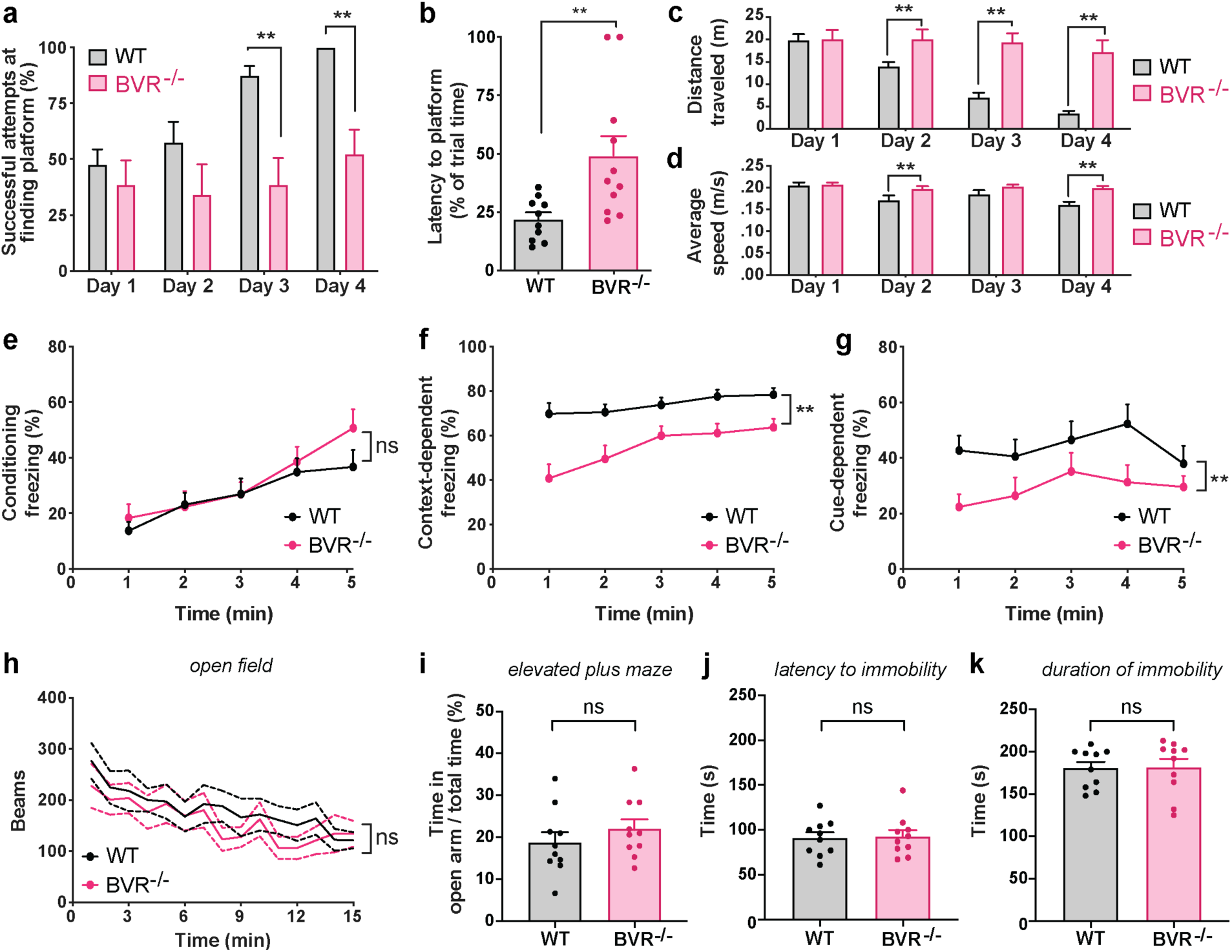
BVR-/- mice exhibit impaired learning and memory, but no anxiety or despair. **a-d**, WT and BVR^-/-^ mice performance in Morris water maze. (*a*) Percent of successful attempts to arrive at platform during trials on days 1-4. (*b*) Latency to arrive at platform on test day 5. (*c-d*) Distance traveled (m) and average swim speed (m/s) during trials on days 1-4. *n* = 10 WT and 11 BVR^-/-^ mice. Points represent individual mice. **e-g**, WT and BVR^-/-^ mice performance in context- and cue- dependent fear conditioning. (*e*) Percent of time that mice remain frozen during training trial in which a series of tone-shock pairs. (*f*) Percent of time that mice remain frozen when re-exposed to training chamber the following day without delivering any shocks. (*g*) Percent of time that mice remain frozen when re-exposed to training tone in a new chamber the following day without delivering any shocks. *n* = 14 WT and 10 BVR^-/-^ mice. Points represent mean of data. **h**, WT and BVR^-/-^ mouse open field locomotor activity over time. *n* = 5 WT and 5 BVR^-/-^ mice. **i**, Percent of time WT and BVR^-/-^ mice open arms of the elevated plus maze. *n* = 10 WT and 10 BVR^-/-^ mice. Points represent individual mice. **j-k**, WT and BVR^-/-^ mice performance in forced swim test. (*j*) Latency to and (*k*) duration of immobility, defined as the absence of movement or paddling with only a hind leg to keep head above water. *n* = 10 WT and 10 BVR^-/-^ mice. Points represent individual mice. (*a*-*g*, *i*-*k*) Mean ± SEM depicted. (*h*) Mean ± 95% CI depicted. *\* P* < 0.05, ** *P* < 0.01, ns = *P* > 0.05, two-tailed unpaired Student’s t-test.

Fear conditioning is an additional form of learning distinct from spatial learning in which organisms learn to predict aversive events. Fear conditioning is a behavioral paradigm in which an offending stimulus becomes associated with an otherwise neutral context or cue. Notably, fear conditioning is thought to be learned, stored, and expressed in both the amygdala and hippocampus through the same principal synaptic signaling that encodes spatial learning (*21*). Electrophysiological studies have demonstrated that neurons in the amygdala undergo LTP (*22, 23*), and pharmacological studies that perturb synaptic signaling impede learning of conditioned responses (*24*). To probe whether BVR also influences fear learning, we compared how well WT and BVR^-/-^ mice associate a context or cue with an aversive sensory stimulus. In these tests, we initially train the mice to freeze in response to a sound by mildly shocking them in the foot shortly afterwards. With each subsequent tone-shock pair, more mice will freeze after the tone in anticipation of being shocked. The following day, the mice are returned to the same chamber as the day before. Mice which associate the chamber with being shocked will freeze, whereas those that do not are more likely to move about. We then move the mice to a new chamber and replay the same sound from the day before. Mice which remember that the sound precedes a shock will freeze in fear, whereas those which do not are again likely to continue moving about. Unlike in the Morris water maze in which BVR^-/-^ mice struggled to learn the location of the hidden platform, BVR^-/-^ mice initially learn to associate the tone with a shock and are just as likely to freeze as WT mice during training (**Fig. 1e**). However, BVR^-/-^ mice do not retain either context- or cue-dependent fear learning the day after. When exposed to either the training chamber context or sound cue, BVR^-/-^ mice freeze less often than WT mice (**Fig. 1f-g**).

Though BVR^-/-^ mice performed comparatively worse in both the Morris water maze and fear conditioning tests, both tests could be confounded if BVR^-/-^ mice are more hyperactive and mobile than WT mice. If BVR^-/-^ mice are hyperactive, they may be more preoccupied by swimming in the Morris water maze than by finding the platform. They may also freeze less in fear conditioning tests even if they remember the chamber or cue. To evaluate whether BVR^-/-^ mice are hyperactive, we monitored their mobility in the open field assay, in which we measure the total distance traveled by mice placed in an open chamber. Whereas hyperactive mice would cover a greater distance, we find that BVR^-/-^ mice are as mobile and active as WT mice (**Fig. 1h**).

We also considered that the Morris water maze and fear conditioning tests could be confounded if BVR^-/-^ mice are more anxious or depressed. However, WT and BVR^-/-^ mice score similarly in tests of anxiety and despair (**Fig. 1i-k**). To measure anxiety, we challenged WT and BVR^-/-^ mice to an elevated plus maze test. In this assay, mice are placed in an elevated plus-shaped apparatus with two open and two enclosed arms and monitored for how long they spend in the open versus closed arms. Higher levels of anxiety are associated with more time spent in the enclosed arms, and classical anxiolytics such as benzodiazepines lead mice to spend more time in the open arms (*25, 26*). To measure despair, we challenged WT and BVR^-/-^ mice to the behavioral despair test (or forced swim test). In this assay, mice are forced to swim in a cylinder filled with water from which they cannot escape and are monitored for how long they attempt to escape before they relent and remain immobile, moving only to keep their heads above water. WT and BVR^-/-^ mice spend a similar proportion of time in the open arm in the elevated plus maze test (**Fig. 1i**), suggesting that BVR^-/-^ mice are not more anxious. WT and BVR^-/-^ mice also forgo escape at a similar rate in the forced swim test (**Fig. 1j-k**), suggesting that BVR^-/-^ mice do not abandon behavioral tests more easily. As a result, it is unlikely that BVR^-/-^ mice perform poorly in the Morris water maze and conditioned fear tests because they are hyperactive, anxious, or depressed, but rather because they suffer learning and memory deficits in multiple domains.

### RNA-sequencing reveals downregulated focal adhesion signaling in BVR^-/-^ hippocampi

Though BVR^-/-^ mice performed comparatively worse in two neurobehavioral assays of learning and memory, neither test immediately reveals the underlying molecular pathology. However, deficits in both the Morris water maze and fear conditioning tests crudely localize to the hippocampus. To identify where in the hippocampus BVR may be expressed, we re-analyzed our published single-cell RNA-sequencing data from hippocampi of WT mice (*27*). We identified 13 distinct cell types within the hippocampus, including populations of proliferative cells, astrocytes, radial glia, intermediate progenitors, oligodendrocyte precursor cells, and inhibitory subpopulations of SST- and VIP-positive interneurons. Hippocampal pyramidal neurons further clustered into distinct genetic populations rooted in basic hippocampal circuitry, including the dentate gyrus, CA1, CA2/CA3, subiculum, and entorhinal cortex (**Fig. 2a**). We find that BVR is ubiquitously expressed across all hippocampal cell populations, though it may be slightly enriched in CA1, CA2, and CA3 pyramidal neurons compared to other cell types (**Fig. 2b-c**).

**Figure 2.**
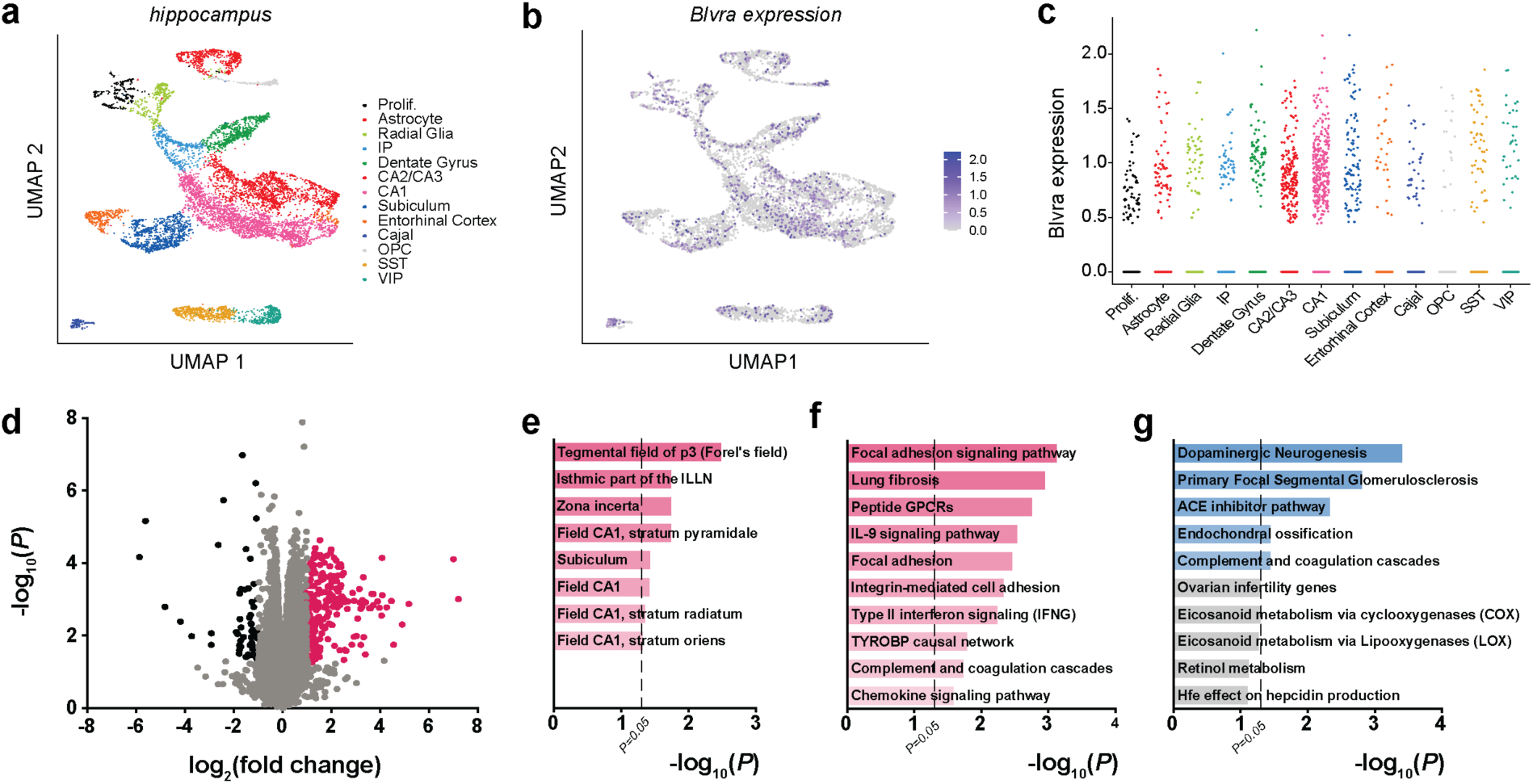
RNA-sequencing reveals downregulated focal adhesion signaling in BVR-/- hippocampi. **a**, UMAP plot of all WT hippocampal cells, colored by cell cluster type. *Prolif*, proliferative cells; *IP*, intermediate progenitors; *Cajal*, Cajal-Retzius cells; *OPC*, oligodendrocyte precursor cells; *SS,* SST+ interneurons; *VIP*, VIP+ interneurons. **b**, Expression of *Blvra* (purple) throughout the hippocampus. **c**, Expression of *Blvra* in each hippocampal cell type. Points represent individual cells. **d**, Volcano plot of transcriptome sequencing of 3 WT and 3 BVR^-/-^ hippocampi. Points represent individual genes. Black points indicate significantly downregulated genes (FDR < 0.5, log_2_(fold change) ≤ -1), whereas pink points indicate significantly upregulated genes (FDR < 0.5, log_2_(fold change) ≥ 1). Gray points are not differentially expressed between genotypes. Notable NRF2 target genes are labeled. **e**, Genetic network analysis of neuroanatomic- and neural cell type- signatures downregulated in BVR^-/-^ hippocampi from transcriptome sequencing in (*d*). **f-g**, Gene ontology analysis of molecular pathways (*f*) downregulated and (*g*) upregulated in transcriptome sequencing in (*d*).

To determine how loss of BVR might affect hippocampal function, we performed a whole-transcriptome RNA sequencing analysis of hippocampi from WT and BVR^-/-^ mice (**Fig. 2d** **and Table S1**). We cross-validated the differential expression profile with quantitative reverse-transcription PCR (qPCR) of several genes (**Fig. S1a-e**). Genetic network analyses detected that loss of BVR is associated with a relative reduction in signature CA1 genes, including those expressed in the stratum pyramidale, stratum radiatum, and stratum oriens (**Fig. 2e** **and Table S2**). BVR^-/-^ hippocampi are also depleted of transcripts linked to the Fields of Forel and zona incerta, discrete components of the subthalamus. Unbiased gene ontology surveys detected that focal adhesion signaling and integrin-mediated cell adhesion are strongly downregulated in BVR^- /-^ hippocampi (**Fig. 2f-g** **and Table S3**). Meanwhile, upregulated genes are more strongly enriched for ontologies associated with dopaminergic neurogenesis, though the rest involve processes with less relevance to the brain such as the nephrotic syndromic primary focal segmental glomerulosclerosis. Several networks of genes involved in complement and chemokine signaling are also dysregulated with loss of BVR (**Fig. 2f-g** **and Table S3**), consistent with a previous study outlining how BVR regulates myeloid chemotaxis in response to complement C5a (*28*).

### Deleting BVR disrupts synaptic focal adhesion signaling in the hippocampus

Comparison of the hippocampal transcriptomes of WT and BVR^-/-^ mice points to the possibility that focal adhesion signaling may be disrupted in the BVR^-/-^ hippocampus. Focal adhesions are large, dynamic structures that bridge the intracellular cytoskeleton to the extracellular matrix (*29*). Integrins span the membrane and anchor focal adhesions by linking extracellular proteins such as fibronectin, laminin, and collagen to actin via adapter proteins such as talin, α-actinin, and vinculin (*30*). Embedded among these more structural proteins is a principal signaling molecule, focal adhesion kinase (FAK). FAK is a non-receptor tyrosine kinase that relays changes in the structure of focal adhesion complexes to major signaling axes within the cell, such as phosphatidylinositol 3-kinase (PI3K) and the mitogen activated protein kinases (MAPKs) (*31*). FAK also translocates to the nucleus and interacts with diverse nuclear factors, including tumor suppressor p53 and NANOG (*32, 33*). As a result, FAK can rapidly and potently transduce extracellular signals into intracellular ones. Though FAK has been most extensively studied in non-neural cells, FAK is also expressed at neural synapses and similarly couples extracellular synaptic signals to long-lasting intracellular changes in synaptic strength (*34–36*). Integrin ligands such as fibronectin and the peptide Gly-Arg-Gly-Asp-Ser-Pro (GRGDSP) stimulate FAK and increase synaptic NMDA and AMPA receptor activity, whereas antibodies against specific integrin subunits prevent the induction of LTP (*37–40*). FAK signaling at the synapse involves much of the same molecular machinery by which it mediates adhesion, motility, and survival in non-neural cells (*41*). In neurons, this machinery also includes the homologous non-receptor tyrosine kinase, proline-rich tyrosine kinase 2 (Pyk2/CAKβ) (*42*).

FAK is classically activated by integrins, which recruit FAK to focal adhesions and induce a conformational change within FAK that favors its activation (*43–45*). FAK can also be activated by G-protein-coupled receptors and by PKC-mediated “inside-out” activation of integrins (*46, 47*). When stimulated, FAK first autophosphorylates itself at Tyr397, generating a high-affinity ligand for the SH2 domain of the kinase Src (*48, 49*). Src then autophosphorylates itself at Tyr416, a crucial regulatory step that leads to a 4-fold increase in Src activity (*50, 51*). Src then reciprocally phosphorylates the catalytic and C-terminal domains of FAK at Tyr407, Tyr576, Tyr577, Tyr871, and Tyr925 (*44*). Like FAK, Pyk2 can be recruited by integrins but is also activated directly by PKC in response to intracellular calcium (*52*). Canonically, active FAK and Pyk2 go on to direct cell migration by activating cytoskeletal proteins such as cofilin and paxillin (*53, 54*). FAK and Pyk2 can also translocate to the nucleus or converge upon parallel PI3K and MAPK signaling (*46, 52, 55*). In neurons, activated Src potently controls synaptic signaling by phosphorylating the NR2B subunit of the NMDA receptor at Tyr1472 (**Fig. 3a**) (*39, 56, 57*). Tyr1472 is basally phosphorylated in the brain but is hyperphosphorylated in CA1 hippocampal slices after tetanic stimulation (*57, 58*), potentiating NMDA conductance (*59–62*) and thereby promoting synaptic plasticity and learning in various paradigms (*37, 39, 63–66*).

**Figure 3.**
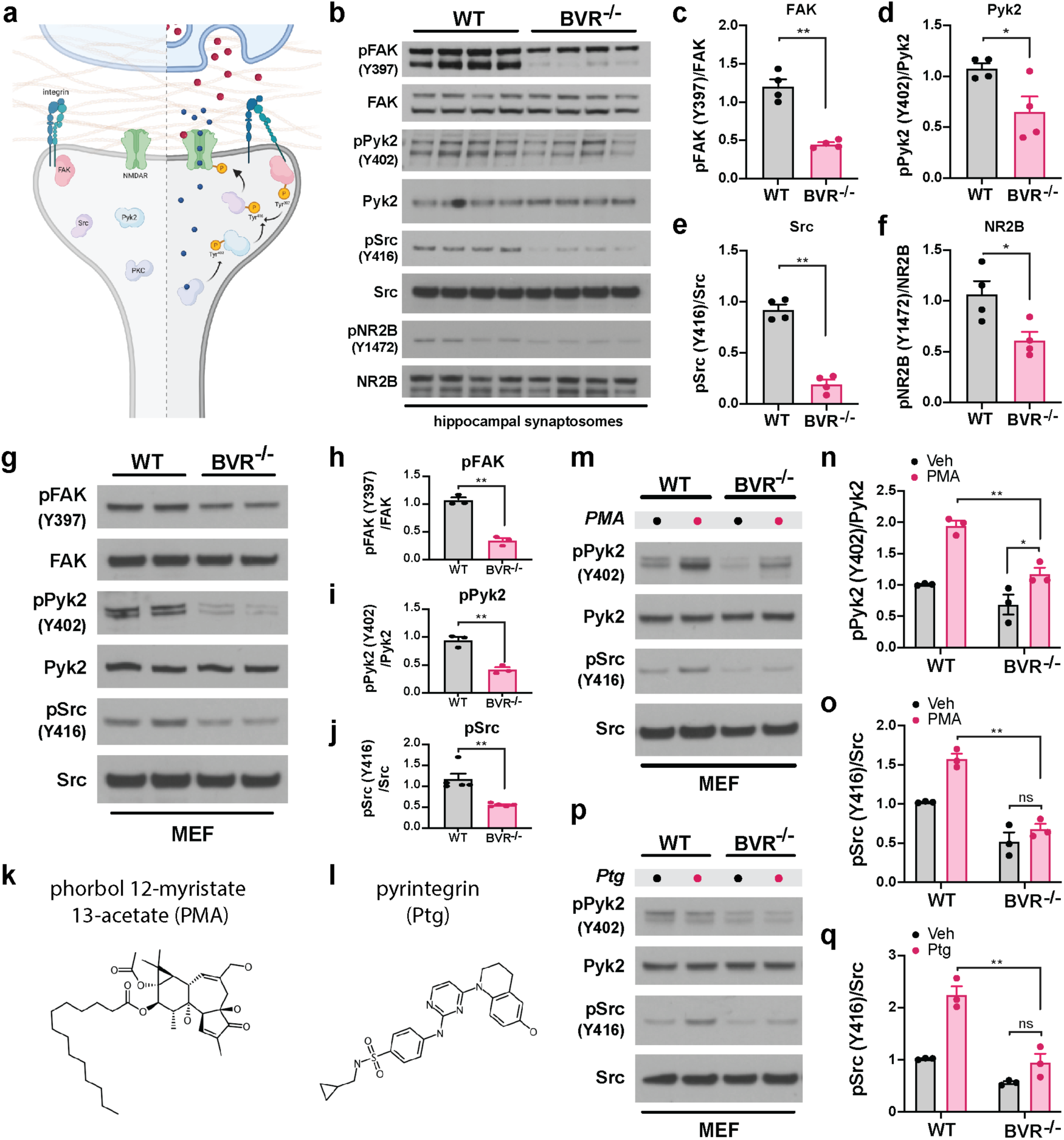
Deleting BVR disrupts synaptic focal adhesion signaling in the hippocampus. **a,** Schematic depicting postsynaptic FAK-mediated signaling pathways at rest (*left*) and during NMDAR-mediated neurotransmission (*right*). Upon glutamate/co-agonist (red circles) binding and depolarization-induced removal of Mg^2+^ block, Ca^2+^ (blue circles) influx through NMDAR activates PKC. Ca^2+^/PKC activate Pyk2 by stimulation of autophosphorylation at Y402 and integrins by direct phosphorylation. The phosphorylation-induced conformational change in integrins stimulates autophosphorylation of FAK at Y397; autophosphorylated FAK and Pyk2 recruit and activate Src by phosphorylation at Y416. Src then binds and phosphorylates the NR2B subunit of the NMDAR at Y1472, an action which potentiates Ca^2+^ influx through the NMDAR. Integrins may also be activated during neurotransmission by secretion of extracellular ligands whose identities are presently unknown. **b-f**, Immunoblots (*b*) and quantifications (*c*-*f*) of phospho-FAK (Y397), FAK, phospho-Pyk2 (Y402), Pyk2, phospho-Src (Y416), Src, phospho-NR2B (Y1472), and NR2B in crude synaptosomes isolated from hippocampi of WT and BVR^-/-^ mice. Data are expressed as normalized ratio of phosphorylated to total protein in each mouse (*n* = 4 per genotype). **g-j**, Immunoblots (*g*) and quantifications (*h*-*j*) of phospho- and total FAK, Pyk2, and Src in WT and BVR^-/-^ MEFs (*n* = 3). **k-l**, molecular structures of (*k*) phorbol 12-myristate 13-acetate (PMA) and (*l*) pyrintegrin (Ptg). **m-q**, Immunoblots (*m*, *p*) and quantification (*n*-*o*, *q*) of phospho- and total Pyk2 and Src in WT and BVR^-/-^ MEFs treated with vehicle and either (*m*-*o*) 500 nM PMA for 30 min or (*p*-*q*) 5 µM Ptg for 5 min (*n* = 3). Points represent data from independent experiments. Mean ± SEM depicted. (*c*-*f*, *h*-*j*) *\* P* < 0.05, ** *P* < 0.01, two-tailed unpaired Student’s t-test. (*n*-*o*, *q*) *\* P* < 0.05, ** *P* < 0.01, ns = *P* > 0.05, two-way ANOVA with *post-hoc* Tukey test.

As hinted at in our whole-transcriptome RNA sequencing analysis, we find that activation of the central components of focal adhesion signaling are profoundly downregulated in BVR^-/-^ hippocampal synaptosomes (**Fig. 3b-f**). The initiatory phosphorylation at Tyr397 in FAK and Tyr402 in Pyk2 is considerably blunted in BVR^-/-^ hippocampal synaptosomes despite equivalent expression of each kinase (**Fig. 3b-d**). Downstream targets of FAK and Pyk2 such as Src at Tyr416 and NR2B at Tyr1472 are hypophosphorylated as well (**Fig. 3b and 3e-f**). When we compare whole extracts from WT and BVR^-/-^ hippocampi however, we do not recover the same pattern we observe in isolated synaptosomes. Neither FAK, Pyk2, Src, nor NR2B are differentially phosphorylated in whole extracts, suggesting that losing BVR impairs synaptic focal adhesion signaling without perturbing the axis throughout the whole cell (**Fig. S2a-e**). Signaling to canonical cytoskeletal regulators downstream of FAK remains intact as well, with no change in the phosphorylation status of p21-activated kinase (PAK) or cofilin (**Fig. S3a-c**). Within synaptosomes, loss of focal adhesion signaling did not bleed into loss of PI3K/Akt or MAPK signaling either (**Fig. S4a-c**).

Since focal adhesion signaling is dynamic and not static, we considered whether losing BVR disrupts acute FAK signaling beyond baseline phosphorylations. In order to acutely manipulate focal adhesion signaling, we isolated and cultured mouse embryonic fibroblasts (MEFs) from WT and BVR^-/-^ mice. Importantly, the same matrix of proteins that is hypophosphorylated in BVR^-/-^ synaptosomes is also dysregulated in BVR^-/-^ MEFs. Tyr397 in FAK, Tyr402 in Pyk2, and Tyr416 in Src are hypophosphorylated in BVR^-/-^ MEFs despite equivalent expression of each kinase (**Fig. 3g-j**). We have reported earlier that neither PI3K/Akt or MAPK signaling is altered in BVR^-/-^ MEFs (*17*).

We induced focal adhesion signaling by two orthogonal approaches in order to resolve where BVR falls within the matrix of proteins involved. Whereas phorbol esters like phorbol 12- myristate-13-acetate (PMA) stimulate PKC to activate FAK/Pyk2 (*67*), integrin ligands such as pyrintegrin act further upstream by engaging integrins (*68*). If neither PMA nor pyrintegrin can recover FAK/Pyk2 phosphorylation in BVR^-/-^ MEFs, BVR may be necessary for activating FAK/Pyk2 themselves. If PMA or pyrintegrin can stimulate FAK/Pyk2 but cannot rescue Src phosphorylation, BVR may instead somehow link active FAK/Pyk2 to Src. In preliminary experiments, we found that Src is predominantly activated by Pyk2 in MEFs and accordingly focused on Pyk2 instead of FAK here.

As expected, PMA activates Pyk2 in WT MEFs with a resulting increase in Src phosphorylation (**Fig. 3m-o**). PMA does increase Pyk2 phosphorylation in BVR^-/-^ MEFs, but the effect is much milder than in WT MEFs. However, PMA does not increase Src phosphorylation, suggesting that stimulating Pyk2 is not sufficient to rescue Src activation without BVR. Acutely applying the integrin ligand pyrintegrin similarly increases Src activation in WT cells but is also unable to overcome the effect of losing BVR in BVR^-/-^ MEFs (**Fig. 3p-q**). As a result, both acute and baseline focal adhesion signaling seem dependent on BVR. BVR appears to be particularly critical for activating Src, since neither integrins nor PKC are able to stimulate Src phosphorylation without BVR. The integrin ligand pyrintegrin does not quantifiably increase in Pyk2 phosphorylation in WT or BVR^-/-^ MEFs, which may be a reflection of how Pyk2 is not as strongly coupled to integrins as FAK (*52, 69*).

### BVR physically interacts with FAK/Pyk2 and Src in a reductase-independent manner

With both acute and baseline focal adhesion signaling dependent on BVR, we considered whether BVR may mediate focal adhesion signaling directly. Since FAK/Pyk2 is unable to activate Src without BVR (**Fig. 3m-q**), we wondered whether BVR may bridge FAK/Pyk2 to Src. If so, BVR might physically associate with FAK, Pyk2, and Src. Consistent with such a model, we find that glutathione-S-transferase (GST)-tagged BVR pulls down FAK, Pyk2, and Src when independently co-expressed in HEK293 cells (**Fig. 4a-c**). GST by itself does not interact with either of the three kinases. We explored whether BVR also interacts with the NMDA receptor subunit NR2B as well, but did not find that the two associate (**Fig. 4d**). To judge the specificity of the interaction between BVR and FAK, we tested whether a related, but structurally dissimilar enzyme also associates with FAK, Pyk2, or Src. Like BVR, the enzyme biliverdin reductase-β (BVRβ) also reduces biliverdin to bilirubin. However, classical BVR (also known as BVRα) and BVRβ reduce biliverdin to entirely different bilirubin isomers and share little-to-no sequence similarity (*70*). We find that BVRβ does not interact with any of the three kinases (**Fig. 4e** **and S5a-b**), suggesting to us that the interactions between FAK, Pyk2, Src, and BVRα are unlikely to be nonspecific. BVR does not interact with an endogenous variant of FAK (FAK-related non kinase or FRNK) either, further indicating that the interactions between BVR and FAK/Pyk2 are isoform-specific (**Fig. S5c**).

**Figure 4.**
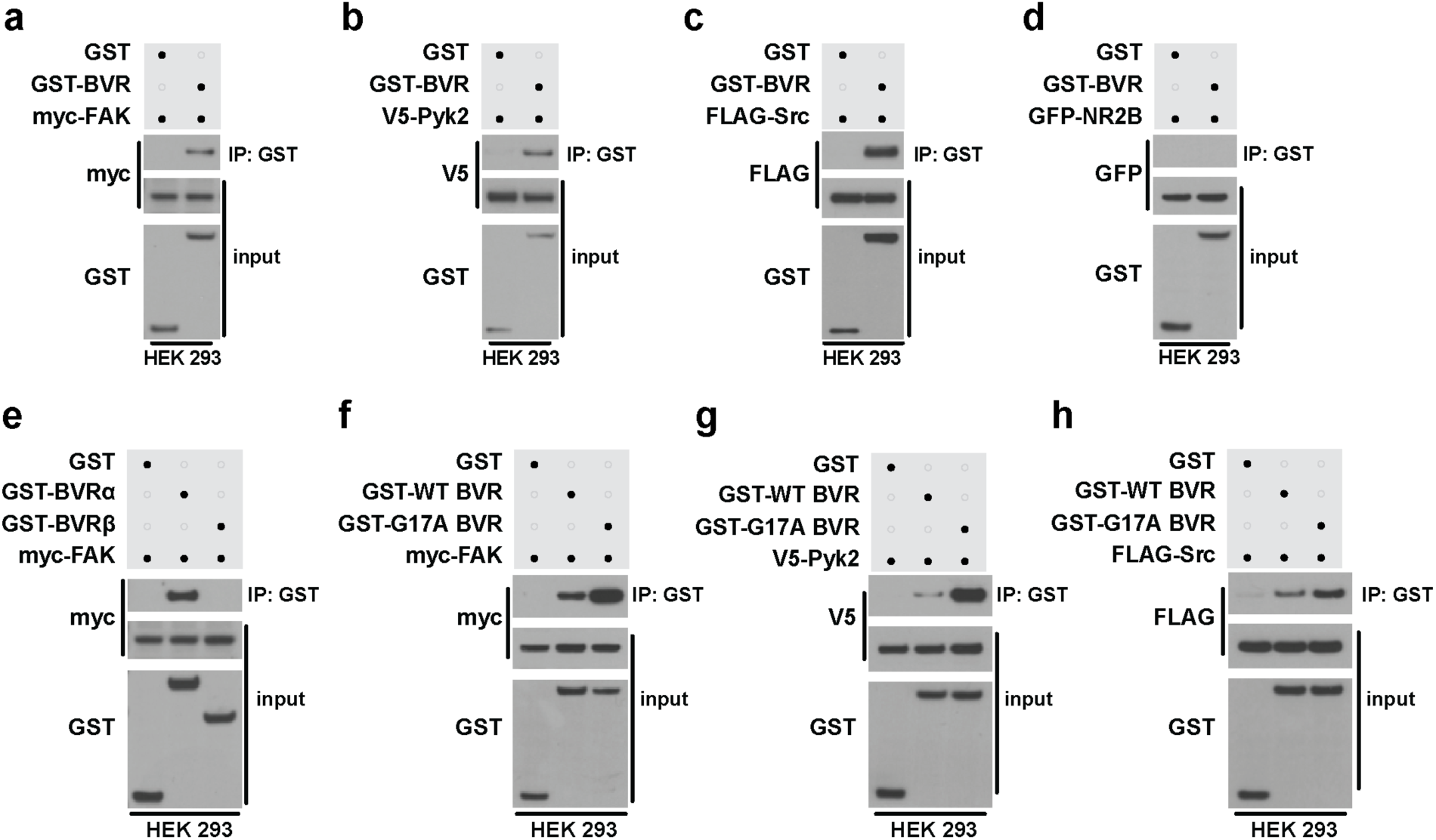
BVR physically interacts with FAK/Pyk2 and Src in a reductase-independent manner. **a-d,** Immunoblots of lysates (input) and GST immunoprecipitates (IP) from HEK 293 cells overexpressing GST or GST-BVR and either (*a*) myc-FAK, (*b*) V5-Pyk2, (*c*) FLAG-Src, or (*d*) GFP-NRF2B. **e,** Immunoblots of lysates (input) and GST IP from HEK 293 cells overexpressing myc-FAK and either GST, GST-BVRα, or GST-BVRβ. **f-h**, Immunoblots of lysates (input) and GST IP from HEK 293 cells overexpressing GST, GST-WT BVR, or GST-G17A BVR and either (*f*) myc-FAK, (*g*) V5-Pyk2, or (*h*) FLAG-Src.

To isolate whether BVR’s canonical oxidoreductase activity is central to its interactions with FAK, Pyk2, or Src, we compared the relative affinity of fully-functional wild-type BVR (WT BVR) for the kinases against a previously characterized catalytically-dead BVR (G17A BVR) (*71*). Crystallographic analyses reveal that Gly17 is key to binding either NADPH or NADH cofactors, and mutating it prevents BVR from reducing biliverdin to bilirubin (*72, 73*). We find that G17A BVR pulls down FAK, Pyk2, and Src to similar or greater degrees than WT BVR, suggesting that the interactions between BVR and FAK/Pyk2 or Src are not dependent on BVR’s enzymatic activity (**Fig. 4f-h**).

### BVR bridges FAK to Src to mediate synaptic focal adhesion signaling

As we sought to confirm these interactions *in vivo*, we found that the tools available to study BVR in the brain were unfortunately inadequate or unreliable. To circumvent these limitations, we generated a mouse line in which BVR is N-terminally tagged with a triplet tandem FLAG octapeptide epitope (Blvra^FLAG^ mice). We introduced the FLAG epitope into the mouse germline just 5’ to the *Blvra* gene using CRISPR/Cas9-directed cleavage and homology-dependent repair (**Fig. 5a**) (*74*). We verified successful insertion of the FLAG epitope into the germline *Blvra* locus by Sanger sequencing, after which we backcrossed the mice over 8 generations to eliminate the possibility of additional, off-target insertions. Western blots from naive Blvra^FLAG^ mice exhibit a single FLAG-immunopositive band between 35-40 kDa, consistent with the predicted molecular weight of 3xFLAG-tagged BVR (**Fig. 5b**).

**Figure 5.**
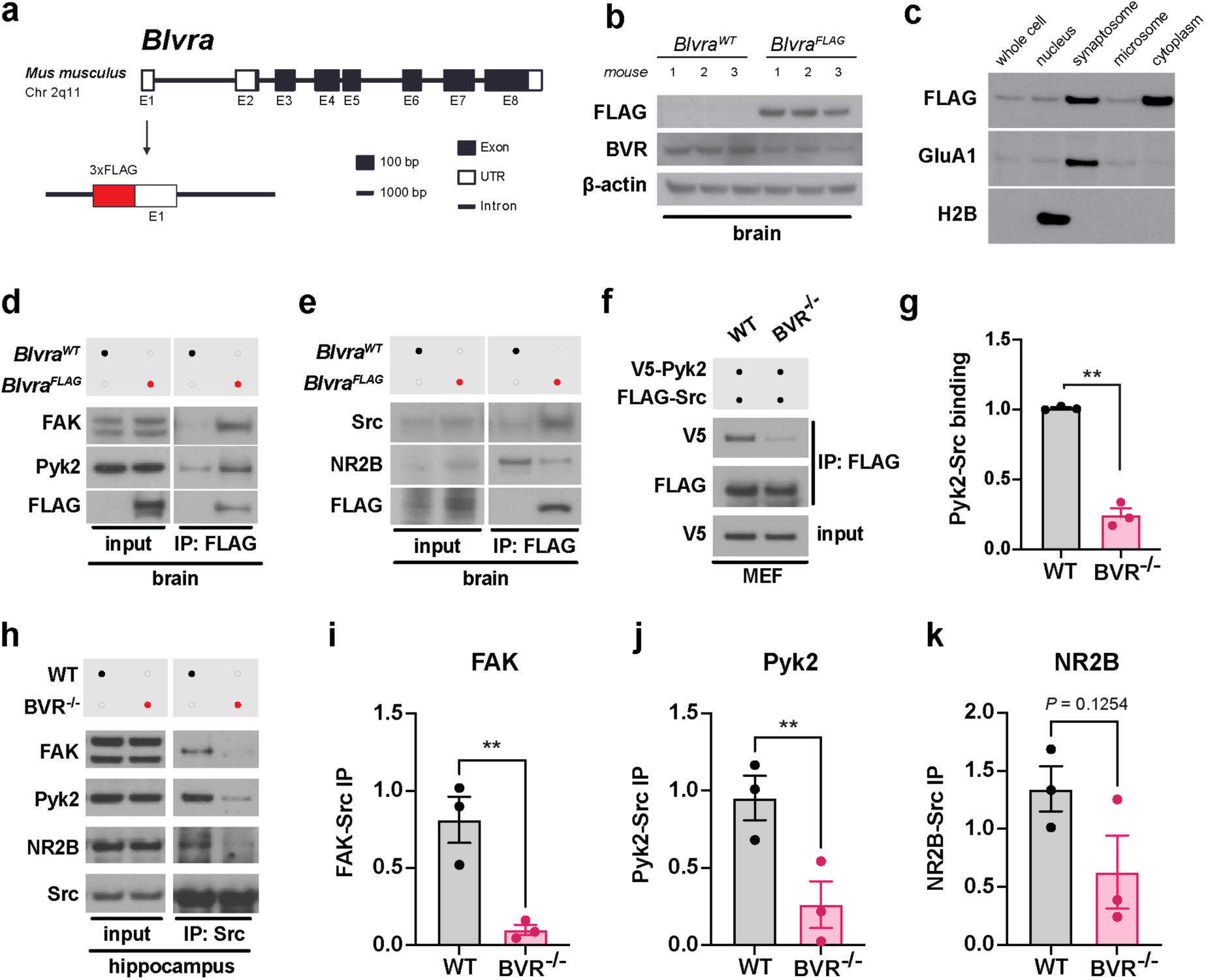
BVR bridges FAK to Src to mediate synaptic focal adhesion signaling. **a**, Schematic of strategy for inserting 3xFLAG tag into the endogenous mouse *Blvra* locus. **b**, Immunoblots of brain lysates from *Blvra^FLAG^* and 3 *Blvra^WT^* mice for FLAG and BVR. β-actin serves as loading control. **c**, Immunoblot of 3xFLAG-BVR in various subcellular fractions of brain tissue from *Blvra^FLAG^* mice. H2B and GluA1 immunoblots were blotted to serve as markers of nuclear and synaptosomal fractions, respectively. **d-e**, Immunoblots of (*d*) FAK, Pyk2, and FLAG-BVR and (*e*) Src, NR2B, and FLAG-BVR in lysates and anti-FLAG immunoprecipitates from *Blvra^WT^* and *Blvra^FLAG^* mouse brain tissue. **f-g**, Immunoblots (*f*) and quantification (*g*) of lysates following anti-FLAG immunoprecipitation (IP) from WT and BVR^-/-^ MEFs overexpressing V5-Pyk2 and FLAG-Src. Data are expressed as normalized ratio of V5-Pyk2 with anti-FLAG IP to FLAG-Src with anti-FLAG IP. V5 input serves as loading control. **h-k**, Immunoblots (*h*) and quantification (*i*-*k*) of lysates following anti-Src IP from WT and BVR^-/-^ hippocampi. Data are expressed as the ratios of (*i*) FAK, (*j*) Pyk2, and (*k*) NR2B that immunoprecipitated with Src, normalized to their respective expression in the input. (*g*, *i*-*k*) Points represent data from independent experiments. Mean ± SEM depicted. ** *P* < 0.01, two-tailed unpaired Student’s t-test.

With BVR successfully tagged, we first leveraged this molecular handle to isolate where BVR may reside within the brain. In fractioning hippocampi from Blvra^FLAG^ mice into nuclear, synaptosomal, microsomal, and cytoplasmic compartments, we find that BVR is present throughout the cell. However, BVR is enriched in synaptosomes (**Fig. 5c**), consistent with a prominent role in synaptic signaling. We then exploited the FLAG epitope to test whether BVR interacts with FAK, Pyk2, and Src *in vivo* in the brain as it does in HEK293 cells (**Fig. 4a-c**). Through a series of experiments in which we immunoprecipitated 3xFLAG-BVR, we find that BVR physically interacts with all three kinases, FAK, Pyk2, and Src (**Fig. 5d-e**). Like in HEK293 cells, BVR does not co-immunoprecipitate with NR2B in the brain.

Since BVR co-immunoprecipitates with both FAK and Src from the brain, we wondered whether BVR may ultimately mediate focal adhesion signaling by physically bridging FAK/Pyk2 to Src. If BVR connects FAK/Pyk2 to Src, FAK/Pyk2 may not be able to bind and stimulate Src without BVR. To explore whether losing BVR leaves FAK/Pyk2 unable to bind Src, we expressed tagged Pyk2 with Src in WT and BVR^-/-^ MEFs and monitored whether less Src co-immunoprecipitates with Pyk2 in BVR^-/-^ MEFs. We observe that less Src co-immunoprecipitates with Pyk2 from BVR^-/-^ MEFs than WT MEFs, suggesting that BVR may indeed bridge FAK/Pyk2 to Src (**Fig. 5f-g**). BVR functions as a fulcrum between FAK/Pyk2 and Src signaling in the hippocampus as well, with Src unable to bind FAK/Pyk2 without BVR. In a series of experiments in which we immunoprecipitated Src, we find that less FAK/Pyk2 co-immunoprecipitates with Src from BVR^-/-^ hippocampi than from WT hippocampi (**Fig. 5h-j**). As a result, Src may interact less with NR2B in BVR^-/-^ hippocampi than in WT hippocampi, though we only observed this in 2 of 3 mice and the difference fell short of statistical significance (*P* = 0.1254) (**Fig. 5h and 5k**).

## Discussion

Synaptic plasticity in the hippocampus is widely considered to be the neurochemical foundation of learning and memory. The NMDA receptor plays a central role by activating a cascade of kinases, enzymes, and second messengers. Following our recent discovery that bilirubin influences NMDA redox signaling (*17*), we considered whether BVR itself may also impact NMDA receptor-dependent synaptic plasticity. In challenging BVR^-/-^ mice with a battery of neurocognitive tasks, we first discover that BVR^-/-^ mice exhibit profound deficits in learning and memory. We uncover that these deficits may be explained by a loss of focal adhesion signaling that is both transcriptionally and biochemically disrupted in BVR^-/-^ hippocampi. Without BVR, FAK, Pyk2, Src, and NR2B are hypophosphorylated and thus presumably hypoactive. We learn that BVR mediates focal adhesion signaling by physically bridging FAK/Pyk2 to Src and that without BVR, FAK/Pyk2 are unable to bind and stimulate Src. Src itself is a molecular hub upon which many signaling pathways converge in order to stimulate the NMDA receptor (*75*), positioning BVR at a prominent intersection of synaptic signaling.

Outside of focal adhesions, BVR is a remarkably pleiotropic protein. Apart from reducing biliverdin to bilirubin, BVR functions as an enzyme-linked receptor that stimulates PI3K/Akt signaling in macrophages (*76*), hepatocytes (*77*), and adipocytes (*78*). BVR also bridges ERK to Elk1 in MAPK signaling in HEK293 cells (*79*). Since we do not observe any difference in the expression or phosphorylation Akt or ERK1/2 between WT and BVR^-/-^ hippocampi however (**Fig. S4a-c**), these alternative functions of BVR may not contribute to its activities in the brain. We have also not observed a difference in the expression, phosphorylation, or nuclear translocation of Akt or ERK1/2 between WT and BVR^-/-^ MEFs previously (*17*). BVR interestingly modulates C5aR1 expression in monocytes (*28*), consistent with our ontology analyses that finds complement and chemokine signaling are dysregulated with loss of BVR (**Fig. 2f-g** **and Table S3**). However, how BVR may influence innate immunity system in the brain remains presently unclear.

While our findings suggest that BVR interfaces with focal adhesions signaling, they do not exclude potential activities of bilirubin or biliverdin. Apart from NMDA superoxide signaling, bilirubin may influence the redox state of other molecules in the synapse. Integrins also interface with redox signaling by triggering oxidation and inhibition of phosphatases that dephosphorylate and inactivate FAK (*80*). As a result, both BVR and FAK are tightly linked to redox signaling. Additionally, loss of BVR in the cortex leads to the accumulation of oxidatively-damaged proteins with impaired autophagy (*81*). Interestingly, shRNA-mediated knockdown of BVR has been shown to impair LTP at CA3-CA1 synapses in hippocampal slices (*82*). Though our data suggest a prominent influence of BVR on hippocampal focal adhesion signaling, FAK and BVR are expressed throughout the brain (*83, 84*), suggesting that BVR may modulate focal adhesion signaling in other brain regions important for memory formation, such as the amygdala and cerebellum.

Importantly, superoxide has been shown to be an essential mediator of LTP and contextual memory (*85, 86*). Thus, while our present data suggest that disruption of BVR impairs learning and memory through dysregulation of FAK- and Src-dependent modulation of the NMDA receptor, it is also possible that loss of bilirubin-mediated superoxide scavenging (*17*) contributes to the learning deficits observed in BVR^-/-^ mice. Further complicating the picture is the fact that superoxide production is necessary for PKC activation during LTP (*85*), suggesting potential crosstalk between redox and integrin-FAK/Pyk2 signaling during synaptic plasticity.

While LTP and the corresponding FAK/Pyk2 activation are generally regarded as NMDA receptor-dependent processes, FAK and Pyk2 can be activated by neurotransmitter receptors other than those for glutamate, including receptors for GABA, acetylcholine, and endocannabinoids (*87–89*). Regulation by BVR of FAK/Pyk2-dependent Src activation may therefore play a role in synaptic signaling mediated by diverse neuromodulators. Bilirubin also binds and activates the nuclear receptor transcription factor peroxisome proliferator-activated receptor alpha (PPAR-α) (*90, 91*), and may thereby also impact synaptic plasticity by altering the composition of the lipid rafts that regulate neurotransmitter exocytosis and postsynaptic receptor clustering (*92, 93*). Along with the mechanisms identified in the present study, these studies may inform future work that discovers how bilirubin and BVR work together to regulate synaptic signaling.

## MATERIALS AND METHODS

### Key Resources Table

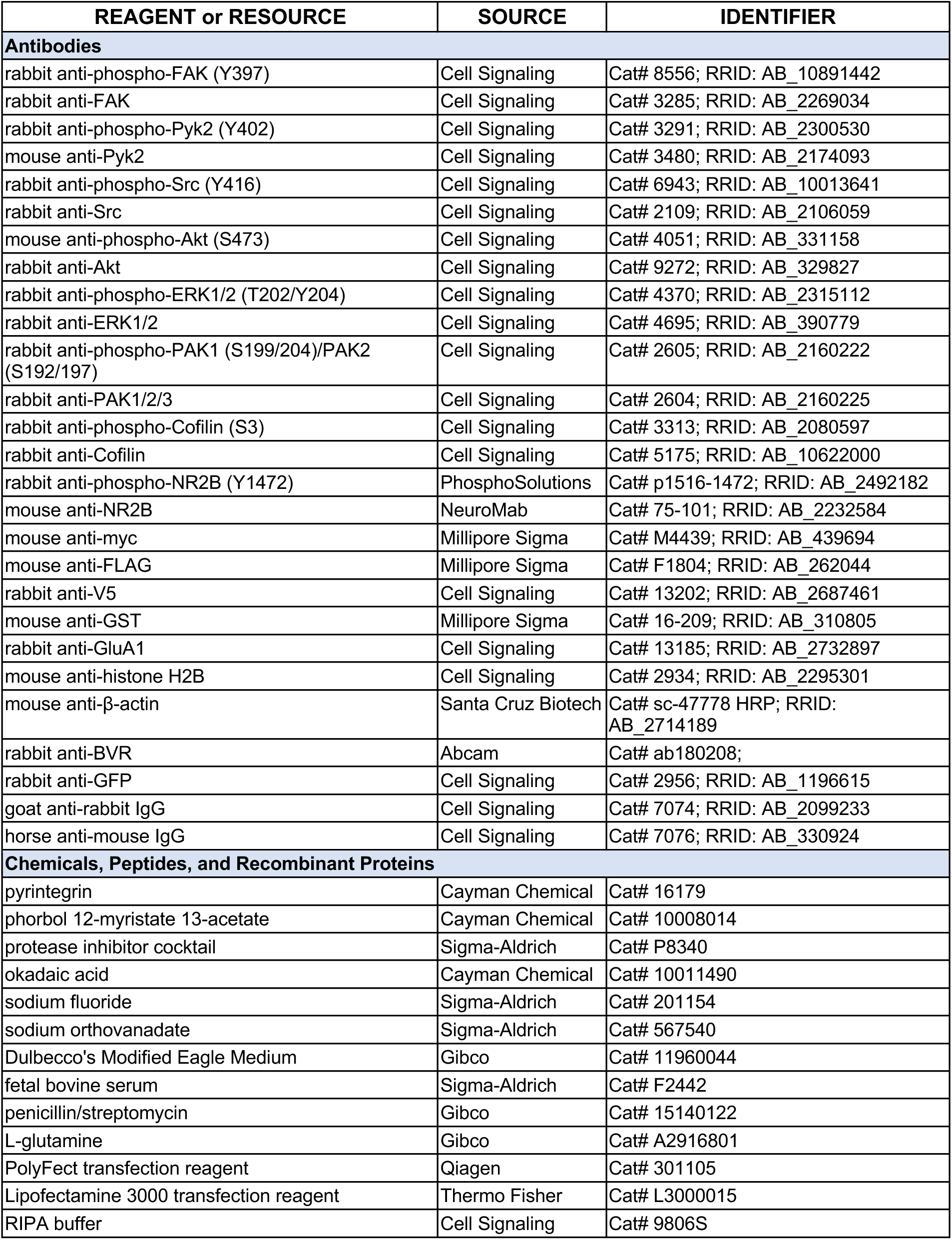
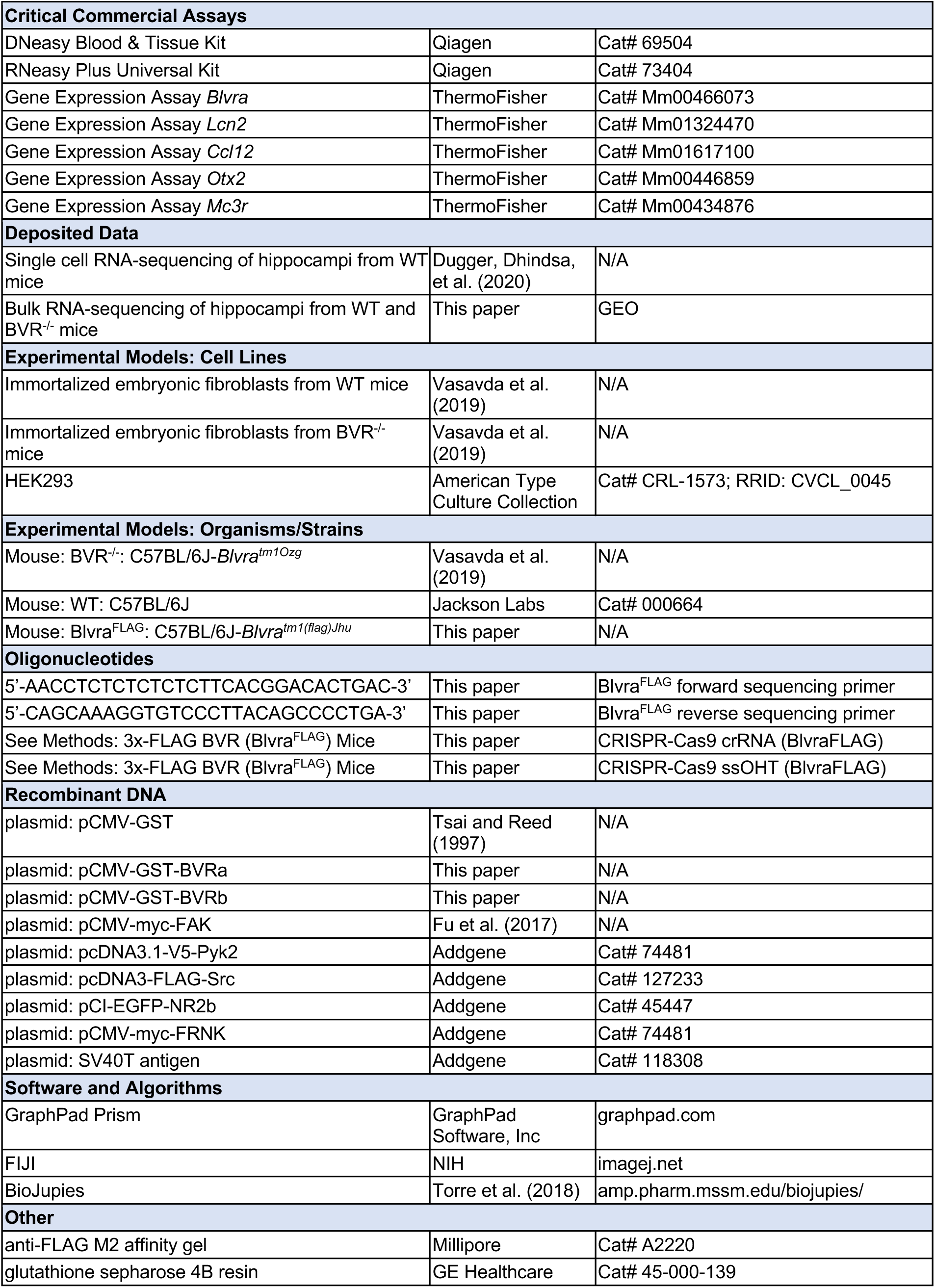

### Lead Contact and Materials Availability

Further information and requests for resources and reagents should be directed to and will be fulfilled by the Lead Contact, Solomon Snyder.

### Experimental Model and Subject Details

#### Mice

##### Animal Care and Use

All experiments were performed in accordance with protocols approved by the Animal Care and Use Committee at the Johns Hopkins University School of Medicine.

##### Wild Type (WT) Mice

C57BL/6J (Stock # 000664) WT mice were purchased from Jackson Labs.

##### BVR Null (BVR^-/-^) Mice

BVR^-/-^ mice were generated as previously described (*17, 94*). Briefly, two LoxP sequences were inserted into the mouse germline flanking exon 3 of *Blvra* for deletion via Cre recombinase. BVR^- /-^ mice were backcrossed to wild-type C57BL/6J mice for fourteen or more generations before these studies. WT littermates were used as controls in behavioral assays. Age-matched wild-type C57BL/6J mice were used as controls for biochemical experiments.

##### 3x-FLAG BVR (*Blvra^FLAG^*) Mice

*Blvra^FLAG^* mice harbor a mutant *Blvra* allele with a triplet tandem FLAG octapeptide epitope (DYKDHD-G-DYKDHD-I-DYKDDDDK) just 5’ of BVR, generating a single 3x-FLAG BVR peptide. This FLAG epitope was inserted into the mouse germline using CRISPR/Cas9-directed cleavage and homology-directed repair (*74*). A CRISPR-Cas9 crRNA (5’- GAGGCCAAGATGAGTACTGAGG-3’) targeting exon 2 of *Blvra* was synthesized by Sigma Aldrich. A single-stranded oligo homology template (5’-ATGGACTACAAAGACCATGACGGTGATTATAAAGATCATGACATCGATTACAAGGATGACGATGACAAGATGAGCACGGAGGTGAGCTGCCCTCAGGGGCTGTAAGGGACACCTTTGCTG-3’) was synthesized by Sigma Aldrich. The gRNA, oligo template, and Cas9 mRNA were injected into C57BL/6 embryos.

Correct insertion of the FLAG epitope was confirmed by Sanger sequencing with the following primers: 5’-AACCTCTCTCTCTCTTCACGGACACTGAC-3’ (Forward) and 5’-CAGCAAAGGTGTCCCTTACAGCCCCTGA-3’ (Reverse). The *Blvra^WT^* allele returns an expected product of 93 nucleotides, whereas the *Blvra^FLAG^* results in a product of 162 nucleotides. Putative *Blvra^FLAG^* mice were then backcrossed to wild-type C57BL/6J mice for eight or more generations before these studies.

### Method Details

#### Materials and Preparation

##### Compounds Were Obtained as Follows

pyrintegrin (Cayman Chemical), phorbol 12-myristate 13-acetate (PMA) (Cayman), sodium fluoride (Sigma-Aldrich), sodium orthovanadate (Sigma-Aldrich), and okadaic acid (Cayman)

##### Material Preparation

All compounds were prepared as 50 μl – 1,000 μl aliquots and stored at -20°C before thawing at 4°C. Freeze/thaw cycles were avoided whenever possible. Pyrintegrin and Phorbol 12-myristate 13-acetate (PMA) were initially dissolved in 100% DMSO and then diluted to 1% DMSO in DMEM.

#### Plasmids/cDNA

Plasmids encoding GST-BVR (GST-BVRα) and GST-BVRβ were generated by cloning cDNA of human *BLVRA* and *BLVRB* into pCMV-GST vector (Clontech/TaKaRa). The construct encoding myc-FAK was generated as previously described (*95*). Plasmids encoding V5-Pyk2 (Addgene Plasmid #127233), FLAG-Src (Addgene Plasmid #118308), GFP-NR2B (Addgene Plasmid # 45447), myc-FRNK (Addgene Plasmid # 74481), and SV40T antigen (Addgene Plasmid #21826) were obtained from Addgene.

#### Pharmacologic Treatments

##### Pyrintegrin and Phorbol 12-myristate 13-acetate (PMA)

Cells were treated with either pyrintegrin or PMA by removing half the volume of culture medium and replacing with an equal volume containing 2x final concentration of drug. Cells were treated with either pyrintegrin (5 µM final, 5 min) or PMA (500 nM final, 30 min).

#### Neurocognitive Behavior Assays

Behavioral tests were performed on 2-4 months old WT and BVR^-/-^ littermate male and female mice. Behavior was assessed concomitantly or in a blocked manner with consideration for both genotype and assay.

##### Morris Water Maze

Spatial learning was assessed with the Morris water maze. In this assay, mice were trained over four days to locate a submerged platform in a pool circular pool (120 cm diameter) filled 45 cm deep with opaque tap water. Each day, we began by revealing the platform to the mice with unique visual cues prominently displayed on the walls of the pool. We then placed the mice at semi-random locations around the pool four times and timed how long it takes them to re-locate the platform as a measure of their spatial learning. Mice were allowed 60 s to reach the platform. After four days of training, the mice are tasked once again with locating the platform but without being shown it beforehand. Each mouse’s location, speed, and latency to find the platform were recorded and automatically scored by EthoVision (Noldus Information Technology). Metrics measured in the Morris water maze were successful attempts at finding the platform, latency to reach the platform, total distance traveled, and swim speed.

##### Fear Conditioning

Fear conditioning was assessed with cue- and context-dependent conditioning tests. The day before training, the mice were first were placed individually in the shock boxes and allowed to acclimatize to the environment for at least 30 min. The following day, mice were placed back in the shock box for training and allowed to reacclimatize for 2 min. A 20 s tone at 80 db and 2000 Hz was delivered, followed by a 2 s 0.5 mA shock to the foot shortly afterwards. Tone–shock pairs were delivered a total of four times over training. The day after, the mice are first returned the same shock chamber. They were monitored for 5 min for freezing behavior. Mice which associate the chamber with being shocked will freeze, whereas those that do not are more likely to move about. We then moved the mice to a new chamber and replay the same 20 s tone from the day before. Mice which remember that the sound precedes a shock will freeze in fear, whereas those which do not are again likely to continue moving about. Freezing was automatically recorded and scored using Cleversys Freezescan (Cleversys).

##### Open Field

Hyperactivity was assessed with the open field test. Baseline activity in the open field was evaluated using activity chambers equipped with infrared beams (San Diego Instruments). Behavioral assays were initiated by placing mice were placed in the center of an enclosed acrylic chamber. Total locomotor activity was automatically recorded and analyzed via photobeams in the x, y, and z axes. Changes in locomotion were considered relative to WT littermate controls.

##### Elevated Plus Maze

Anxiety was assessed with the elevate plus maze test. In this assay, mice were placed in a plus-shaped apparatus 1 m above the floor, built of two open and two enclosed runways pointing in orthogonal directions (San Diego Instruments). Mice were monitored for how long they spend in the open versus closed runways. Tests were initiated by placing mice in the center of the maze, which is neither in the open arms nor the closed arms. Each challenge lasted 5 min.

##### Forced Swim

Despair was assessed with the forced swim test. In this assay, mice are challenged with two brief trials during which they are forced to swim in a 15 x 30 cm glass cylinder filled with room temperature water from which they cannot escape. During the second trial, mice are monitored over the course of 6 min for how long they attempt to escape before they relent and remain immobile. Immobility is defined as the absence of movement or paddling with only a hind leg to keep their heads above water.

#### Western Blotting

Western blotting was performed as previously described (*17*). Briefly, cells and tissues were homogenized at 4°C in lysis buffer (pH 7.4 solution of 50 mM Tris-HCl, 150 mM NaCl, 0.1% SDS, 0.5% sodium deoxycholate, and 1% Triton X-100) supplemented with protease inhibitors (Sigma), 50 mM sodium fluoride, 200 µM sodium orthovanadate, and 125 nM okadaic acid (Cayman). Lysates were then pulse sonicated and centrifuged at 16,000*g* for 10 min at 4°C. Fifteen micrograms of cleared lysate were run on a 4-12% polyacrylamide Bis-Tris gradient gel in running buffer (pH 7.3 solution of 50 mM MES, 50 mM Tris Base, 0.1% SDS, 1 mM EDTA) and then transferred to a PVDF membrane. Membranes were blocked with 5% bovine serum albumin (BSA) (w/v) in TBS-T (pH 7.6 solution of 16 mM Tris-HCl, 140 mM NaCl, 0.1% Tween-20) for 1 h at 25°C and then incubated with primary antibodies in 5% bovine serum albumin (BSA) (w/v) in TBS-T overnight at 4°C. The following day, membranes were washed with TBS-T, and then incubated with secondary antibodies in 5% BSA (w/v) in TBS-T for 1 h at 25°C. Protein bands were visualized using SuperSignal West Pico Plus Chemiluminescent Substrate (ThermoFisher). Unless otherwise indicated, blots shown in figures are representative of at least three independent experiments.

The following primary antibodies were used: rabbit anti-phospho-FAK (Y397) (Cell Signaling 8556, 1:1000), rabbit anti-FAK (Cell Signaling 3285, 1:1000), rabbit anti-phospho-Pyk2 (Y402) (Cell Signaling 3291, 1:1000), mouse anti-Pyk2 (Cell Signaling 3480, 1:2000), rabbit anti-phospho-Src (Y416) (Cell Signaling 6943, 1:1000), rabbit anti-Src (Cell Signaling, 2109 1:2000), mouse anti-phospho-Akt (S473) (Cell Signaling 4051, 1:1000), rabbit anti-Akt (Cell Signaling 9272, 1:1000), rabbit anti-phospho-ERK1/2 (T202/Y204) (Cell Signaling 4370, 1:2000), rabbit anti-ERK1/2 (Cell Signaling 4695, 1:2000), rabbit anti-phospho-PAK1 (S199/S204) / PAK2 (S192/S197) (Cell Signaling 2605, 1:1000), rabbit anti-PAK1/2/3 (Cell Signaling 2604, 1:1000), rabbit anti-phospho-Cofilin (S3) (Cell Signaling 3313, 1:1000), rabbit anti-Cofilin (Cell Signaling 5175, 1:1000), rabbit anti-phospho-NR2B (PhosphoSolutions P1516-1472, 1:1000), mouse anti-NR2B (NeuroMab 75- 101, 1:1000), rabbit anti-GluA1 (Cell Signaling 13185, 1:1000), mouse anti-histone H2B (Cell Signaling 2934, 1:1000), rabbit anti-V5 (Cell Signaling 13202, 1:1000), mouse anti-FLAG (Millipore Sigma F1804, 1:2000), mouse anti-myc (Millipore Sigma M4439, 1:4000), mouse anti-GST (Millipore Sigma 16-209, 1:10,000), mouse anti-β-actin (Santa Cruz Biotech sc-47778 HRP; 1:10,000), rabbit anti-BVR (Abcam ab180208, 1:1000), and rabbit anti-GFP (Cell Signaling 2956, 1:1000).

The following HRP-linked secondary antibodies were used: goat anti-rabbit IgG (Cell Signaling 7074; 1:5000) and horse anti-mouse IgG (Cell Signaling 7076; 1:5000).

#### Subcellular Fractionation

Hippocampi were fractionated into nuclear, synaptosomal, microsomal, mitochondrial, and cytoplasmic compartments as previously described (*96, 97*). Briefly, hippocampi were first lysed in 20 volumes of homogenizing buffer (pH 7.3 solution of 0.32 M sucrose and 0.005 M Na_3_PO_4_) at 4°C supplemented with protease inhibitors (Sigma), 50 mM sodium fluoride, 200 µM sodium orthovanadate, and 125 nM okadaic acid (Cayman). Homogenates were then centrifuged at 1,000*g* for 10 min at 4°C, resulting in pellet P1. The supernatant was recentrifuged at 17,500*g* for 20 min at 4°C. The resulting pellet contains a mixture of myelin, synaptosomes, and mitochondria (pellet P2). The supernatant was centrifuged once more at 100,000*g* for 120 min at 4°C to pellet microsomes. The resulting supernatant was considered the cytoplasmic compartment. Pellet P2 was resuspended in homogenizing buffer and subfractioned on a discontinuous sucrose gradient of 0.4, 0.6, 0.8, 1.0, and 1.2 M by centrifugation at 61,000*g* for 120 min at 4°C. Fractions between 0.32-0.8 M sucrose contain myelin and synaptosomal ghosts as evidenced by electron microscopic examination of these fractions. Fractions between 0.8-1.2 M sucrose contain synaptosomes as evidenced by high [^3^H]naloxone and [^3^H]]dihydromorphine binding in previous studies (*98, 99*). Fractions between at 1.2 M sucrose contain mitochondria. Nuclei were purified from pellet P1 by resuspending P1 in a pH 7.4 solution of 1.5 M sucrose, 50 mM Tris-HCl, 5 mM MgCl2, and 25 mM KCl, layering it upon on a solution of 1.8 M, and centrifuging at 40,000*g* for 60 min at 4°C.

#### Immunoprecipitation

Cells or tissue were harvested on ice in lysis buffer (pH 7.6 solution of 50 mM Tris-HCl, 150 mM NaCl, 1% Triton X-100, and 5% glycerol (v/v)) supplemented with protease inhibitor cocktail, passed 10 times through a 27G syringe, and centrifuged at 16,000*g* for 10 min at 4°C. Glutathione Sepharose 4B (GE Healthcare) or anti-FLAG M2 (Sigma) beads were added to cleared lysates and rotated overnight at 4 °C. 5% of each sample was set aside as input prior to addition of beads. The following day, beads were pelleted by centrifugation at 1,000*g* for 1 min at 4 °C. The supernatant was aspirated and beads were washed 5 times in lysis buffer by centrifugation, aspiration, and resuspension. Proteins were eluted from beads by boiling for 5 min in LDS sample buffer (ThermoFisher) and the eluates analyzed by Western blot as described above.

#### Cell Culture

##### Mouse Embryonic Fibroblasts (MEFs)

WT and BVR^-/-^ were isolated as previously described (*17*). Briefly, embryonic day 14.5 (E14.5) embryos were obtained from timed BVR^+/−^ matings. The pups were decapitated and eviscerated, after which the remaining portion was trypsinized and sheared. Isolated MEFs were plated in 2 wells of a 6-well plate and cultured overnight in DMEM (Gibco) supplemented with 10% FBS, 100 U/mL penicillin and streptomycin, and 2 mM glutamine at 37°C with 5% CO_2_. MEFs were then expanded to 6-well plates and transiently transfected with SV40T antigen (Addgene) and maintained until stably proliferative. DNA was isolated from the heads of the pups and genotyped to confirm the genotype of the corresponding MEF cell line.

##### Human Embryonic Kidney (HEK) 293 Cells

HEK293 cells were obtained from the American Type Culture Collection. HEK293 cells were cultured in Dulbecco’s Modified Eagle Medium (DMEM), 10% fetal bovine serum, penicillin/streptomycin (100 U/ml), and glutamine (2 mM) in an atmosphere of 5% CO_2_ at 37°C.

##### Transfection

HEK293 cells and MEFs were transiently transfected with PolyFect (Qiagen) and Lipofectamine 3000 (Thermo Fisher) transfection reagents, respectively, according to manufacturer’s instructions. Cells were harvested 48-72 hours after transfection.

#### RNA-sequencing and Analysis

##### Single-Cell RNA-Sequencing

Single-cell RNA-sequencing was performed as previously described (*27*). Briefly, hippocampi from postnatal day 0 pups and dissected and dissociated for sequencing. Single cell RNA-seq libraries were constructed using the 10X Chromium Single Cell 3′ Reagent Kits v2 according to manufacturer descriptions and subsequently sequenced on a NovaSeq 6000. Reads were aligned to the mm10 genome using the 10X CellRanger pipeline with default parameters to generate the feature-barcode matrix.

Downstream quality control and analyses on feature-barcode matricies were performed with Seurat v3. Genes that were not detected in at least 4 cells were excluded. Cells with fewer than 1,000 genes or more than 5,000 genes were also excluded from analysis. We also excluded cells in which greater than 15% of reads mapped to mitochondrial genes. The filtered matrices were log-normalized and scaled to 10,000 transcripts per cell.

Using the variance-stabilizing transformation in the FindVariableFeatures function, we identified the top 2,000 most variable genes per sample. We then harmonized gene expression across datasets prior to clustering by identifying anchors between samples in each dataset using the FindIntegrationAnchors function and then computing an integrated expression matrix from these anchors as input to the IntegrateData function. We then linearly regressed the number of UMIs per cell and percentage of mitochondrial reads using the ScaleData function and performed dimensionality reduction using PCA. For each dataset, we selected the top 30 dimensions to compute a cellular distance matrix, which was used to generate a K-nearest neighbor graph. The KNN was used as input to the Louvain Clustering algorithm implemented in the FindClusters function. We chose a resolution parameter of 0.8 for clustering via Louvain. To annotate and merge clusters, we performed differential gene expression analysis on the integrated expression values between each cluster using the default parameters in the FindMarkers function.

##### Bulk RNA-Sequencing

We performed bulk RNA-sequencing to profile the differential transcriptional signatures of WT and BVR^-/-^ hippocampi. Total RNA was isolated from three WT and three BVR-/- hippocampi from four-month-old male mice using TRizol and RNeasy Plus Universal Kit (Qiagen) as per the manufacturer’s instructions. DNA was digested with DNase treatment. mRNA was enriched by poly-A selection and prepped for paired-end reads using an Illumina TruSeq mRNA sample preparation kit with 100-base pair adapters. Samples were sequenced by Illumina Novaseq.

Differential gene expression analysis was performed using BioJupies (*100*). We leveraged BioJupies to perform Pathway Enrichment Analyses through Enrichr (*101*) to identify the biological processes that are over-represented in the gene sets up-regulated and down-regulated by in BVR^-/-^ hippocampi.

#### DNA/RNA Isolation, PCR, and Quantitative-PCR (q-PCR)

Total cellular or tissue DNA/RNA was extracted using the DNeasy Blood & Tissue Kit (Qiagen) for DNA or RNeasy Plus Universal Kit (Qiagen) for RNA per the manufacturer’s instructions. PCR was performed with the Platinum Taq DNA Polymerase High Fidelity Kit (Invitrogen).

### Quantification and Statistical Analysis

All data were plotted and expressed as the median and range, mean ± SEM, or mean ± 95% CI, as noted. Statistical comparisons were performed using two-tailed unpaired Student’s *t*-tests, Fisher’s exact tests, or ANOVA analyses, as noted. Differences were considered significant at *P* < 0.05. All *in vivo* experiments were performed concomitantly or in a blocked manner with consideration for both genotype and treatment.

### Data and Code Availability

All data generated or analyzed during this study are included in this article.

## Supporting information

BVR-NMDA-Vasavda et al-Supplemental Material

BVR-NMDA-Vasavda et al-TableS1

BVR-NMDA-Vasavda et al-TableS2

BVR-NMDA-Vasavda et al-TableS3

## ACKNOWLEDGMENTS

This work was supported by NIH NIDA grant P50 DA044123 (to S.H.S.), T32 GM136577 (to C.V.), and Research Scholar Award from the Neurosurgery Pain Research Institute at Johns Hopkins Medicine (to C.V.). Illustrations were created with BioRender.

## AUTHOR CONTRIBUTIONS

C.V. conceptualized the work. C.V. and E.R.S. designed the experiments. C.V., E.R.S., J.L., R.K., R.S.D., S.S., C.R., R.T., A.M.S., L.A., and B.D.P. guided or performed the experiments. C.V., E.R.S., R.K., R.S.D., and R.T. analyzed the data. C.V., E.R.S., and S.H.S. wrote the manuscript. All authors reviewed and edited the manuscript.

## DECLARATION OF INTERESTS

The authors declare no competing financial interests.

Table S1. Differential gene expression between WT and BVR-/- hippocampi.

Table S2. Genetic network analysis of neuroanatomic- and neural cell type- signatures downregulated in BVR-/- hippocampi.

Table S3. Gene ontology analysis of differential gene expression between WT and BVR-/- hippocampi.

---

**Figure S1.**
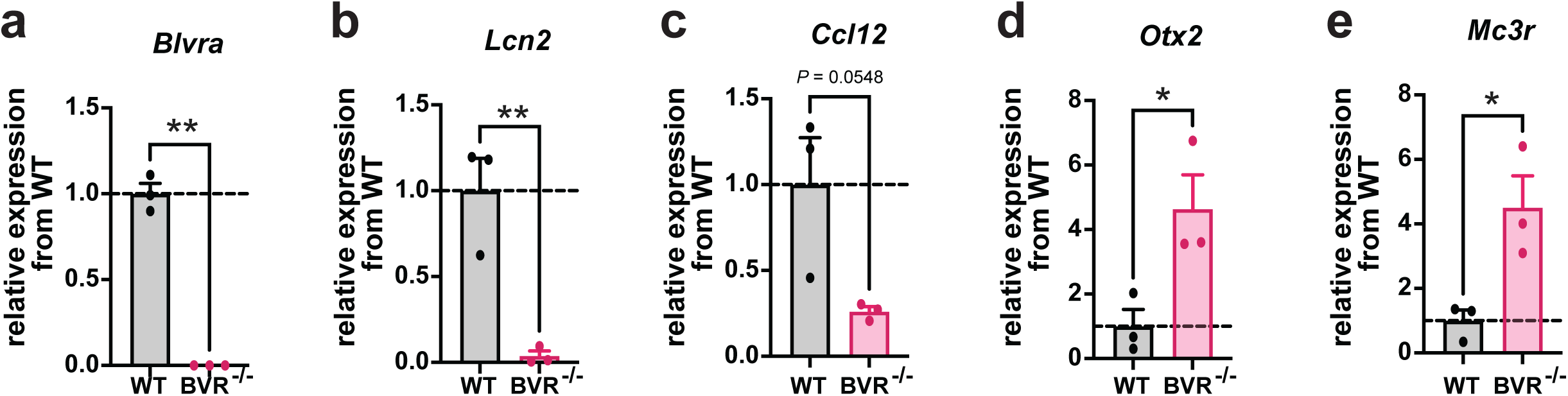
Validation of RNA sequencing from WT and BVR-/- hippocampi. **a-d**, Quantitative PCR analysis of *Blvra*, *Lcn2*, *Ccl12*, *Otx2*, and *Mc3r* mRNA from WT and BVR^-/-^ hippocampi, normalized to β-actin. Points represent individual mice. Mean ± SEM depicted. *\* P* < 0.05, ** *P* < 0.01, two-tailed unpaired Student’s t-test.

**Figure S2.**
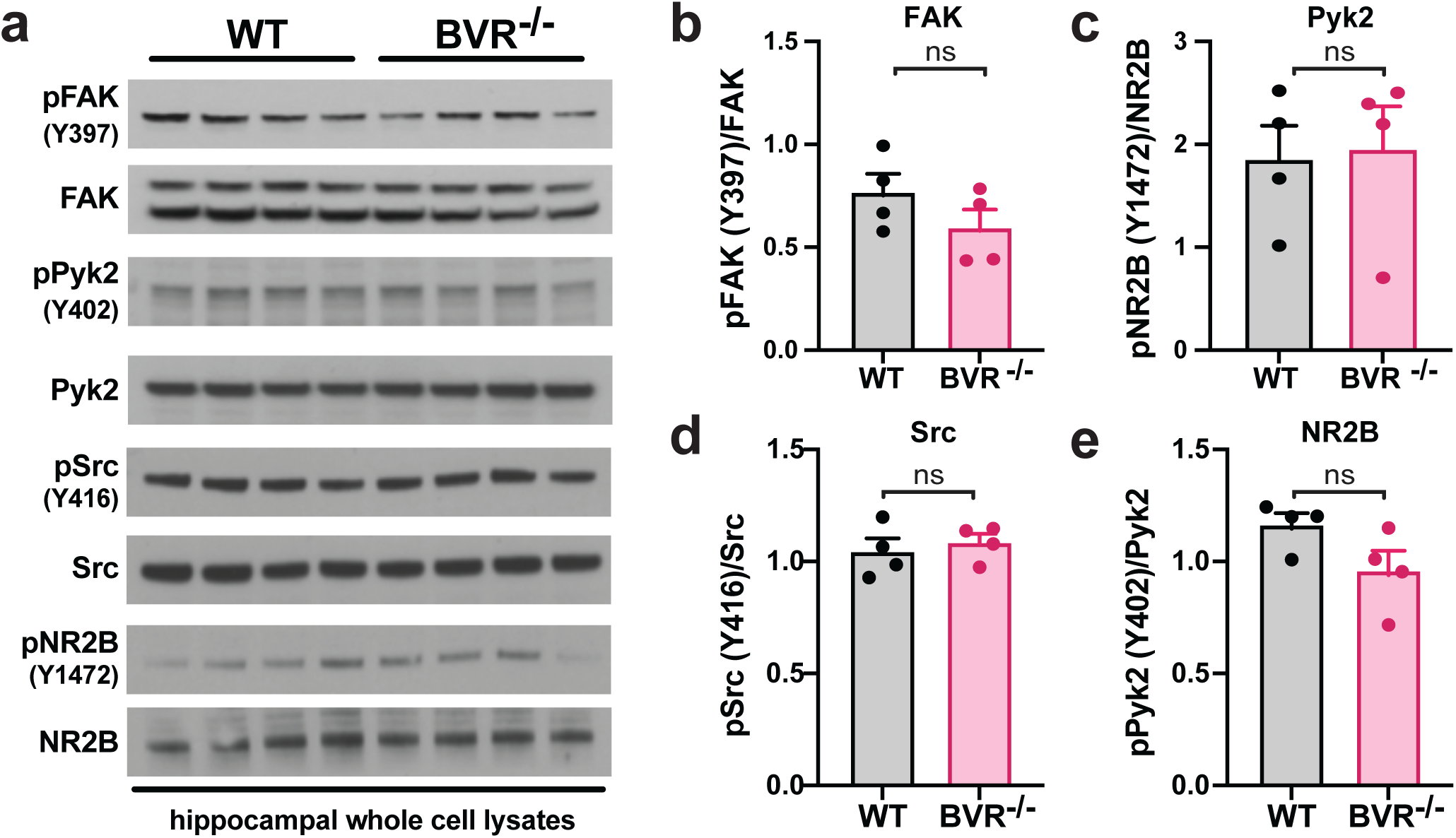
Focal adhesion signaling is not globally disrupted in BVR-/- hippocampi. **a-e**, Immunoblots (*a*) and quantifications (*b*-*e*) of phospho-FAK (Y397), FAK, phospho-Pyk2 (Y402), Pyk2, phospho-Src (Y416), Src, phospho-NR2B (Y1472), and NR2B in hippocampal whole cell lysates from WT and BVR^-/-^ mice. Data are expressed as normalized ratio of phosphorylated to total protein in each mouse. Points represent data from independent experiments. Mean ± SEM depicted. ns = *P* > 0.05, two-tailed unpaired Student’s t-test.

**Figure S3.**
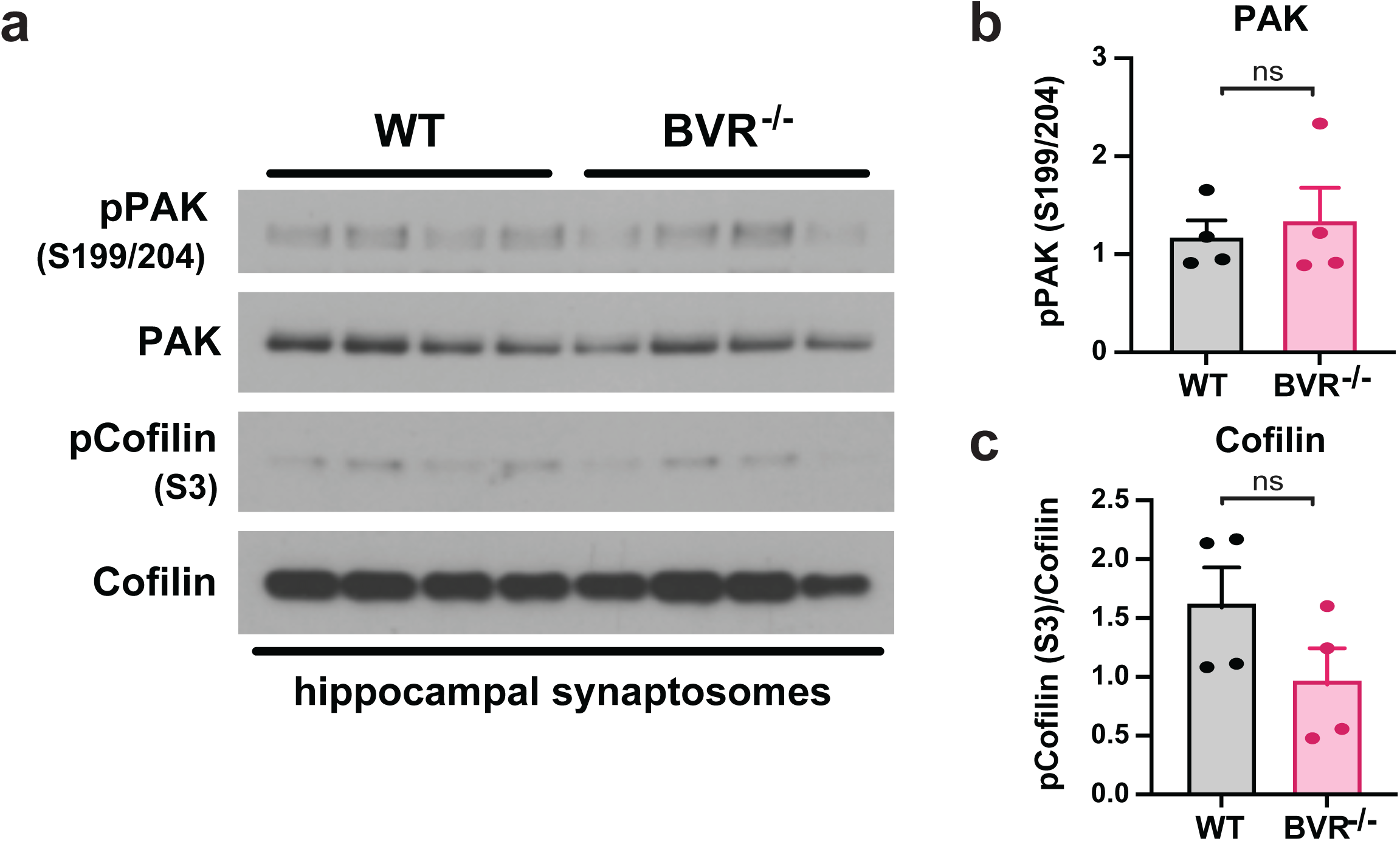
Canonical FAK signaling is intact in BVR-/- synaptosomes. **a-c**, Immunoblots (*a*) and quantifications (*b*-*c*) of phospho-PAK (S199/204), PAK, phospho-Cofilin (S3), and Cofilin in crude synaptosomes isolated from hippocampi of WT and BVR^-/-^ mice. Data are expressed as normalized ratio of phosphorylated to total protein in each mouse. Points represent data from independent experiments. Mean ± SEM depicted. ns = *P* > 0.05, two-tailed unpaired Student’s t-test.

**Figure S4.**
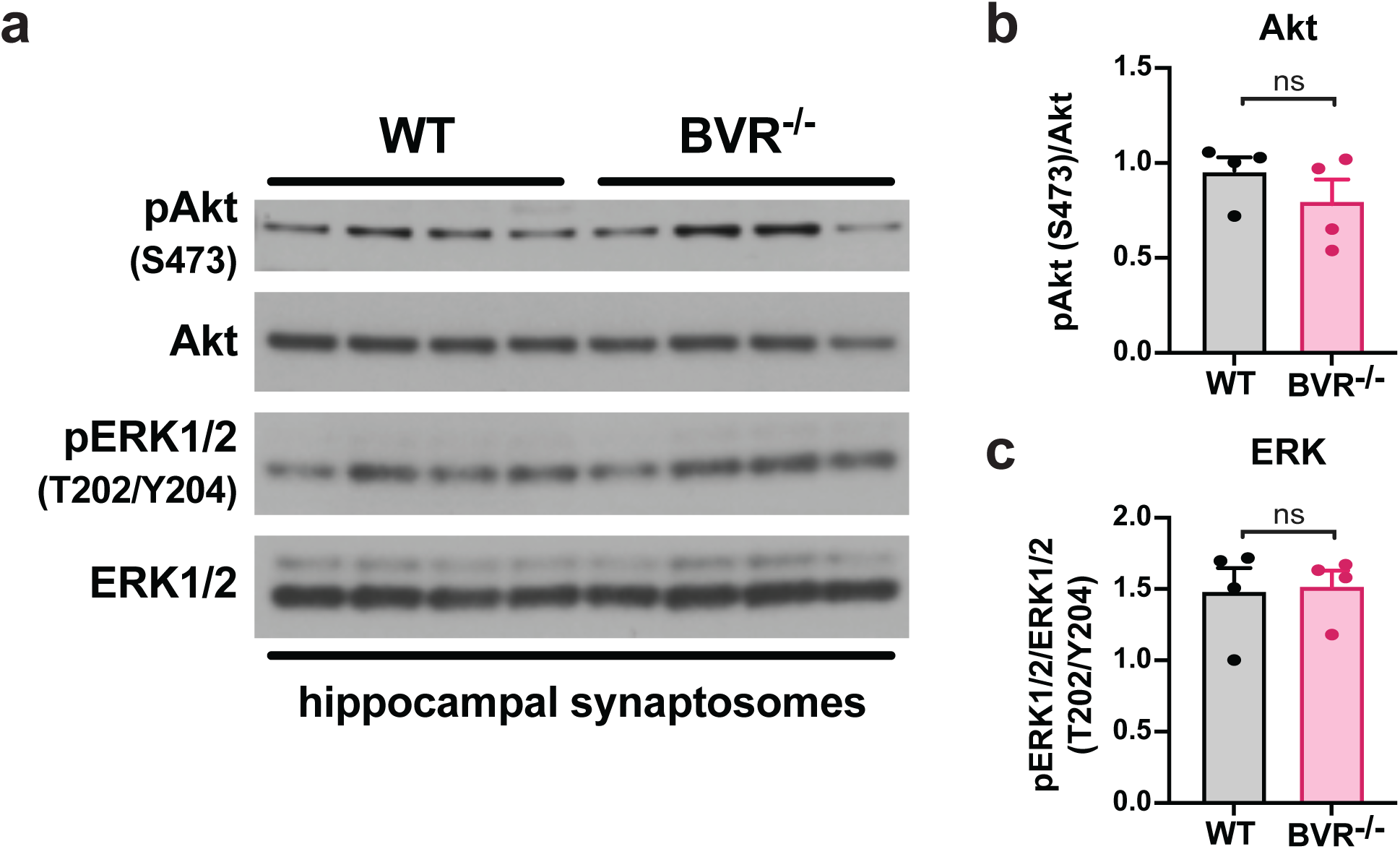
PI3K/Akt and MAPK signaling is not disrupted in BVR^-/-^ hippocampal synaptosomes. **a-c**, Immunoblots (*a*) and quantifications (*b*-*c*) of phospho-Akt (S473), Akt, phospho-ERK1/2 (T202/Y204), and ERK1/2 in crude synaptosomes isolated from hippocampi of WT and BVR^-/-^ mice. Data are expressed as normalized ratio of phosphorylated to total protein in each mouse. Points represent data from independent experiments. Mean ± SEM depicted. ns = *P* > 0.05, two-tailed unpaired Student’s t-test.

**Figure S5.**
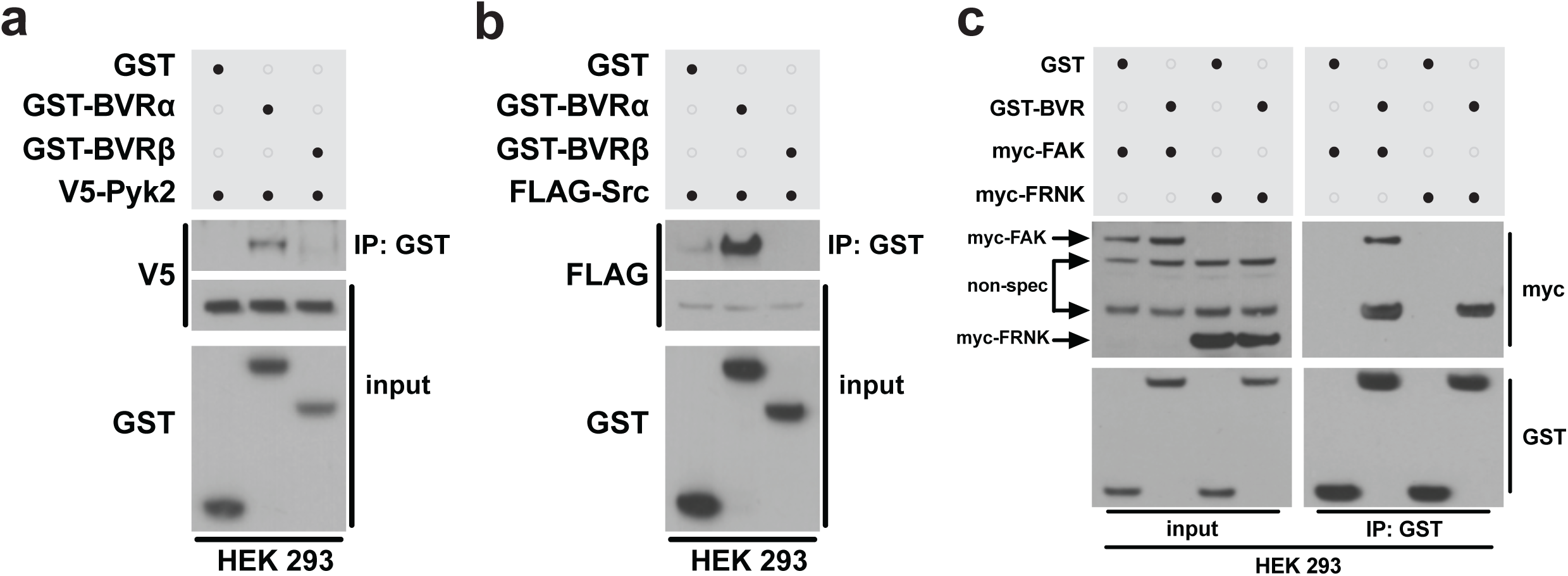
BVR interaction with FAKs is isoform-specific. **a-b**, Immunoblots of lysates (input) and GST immunoprecipitates (IP) from HEK 293 cells overexpressing (*a*) V5-Pyk2 or (*b*) FLAG-Src and either GST, GST-BVRα, or GST-BVRβ. **c**, Immunoblots of lysates (input) and GST IP from HEK 293 cells overexpressing GST or GST-BVR and either myc-FAK or myc-FRNK. Immunoreactive bands corresponding to myc-FAK, myc- FRNK, or a nonspecific signal are indicated by labeled arrows.

## Notes

### Competing Interest Statement

The authors have declared no competing interest.

### Summary of Updates

Additional results included.

