## Supplementary material for "Biliverdin reductase bridges focal adhesion kinase to Src to modulate synaptic signaling": BVR-NMDA-Vasavda et al-Supplemental Material

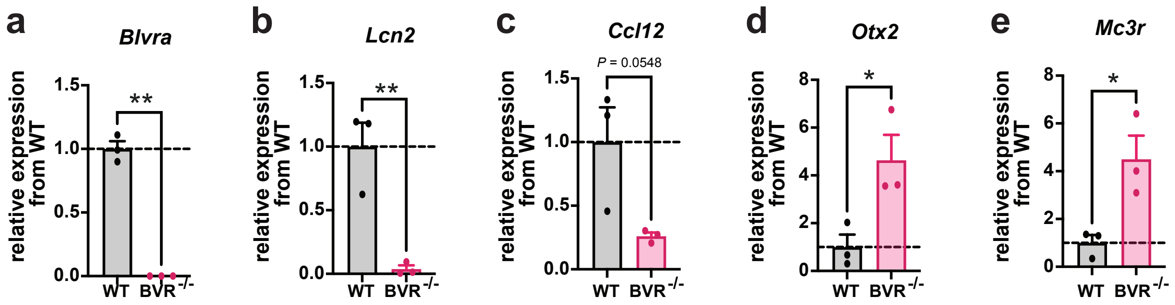


**Figure S1. Validation of RNA sequencing from WT and BVR-/- hippocampi.**

**a-e**, Quantitative PCR analysis of *Blvra*, *Lcn2*, *Ccl12*, *Otx2*, and *Mc3r* mRNA from WT and BVR$^{-/-}$ hippocampi, normalized to β-actin. Points represent individual mice. Mean ± SEM depicted. * *P* < 0.05, ** *P* < 0.01, two-tailed unpaired Student's t-test.

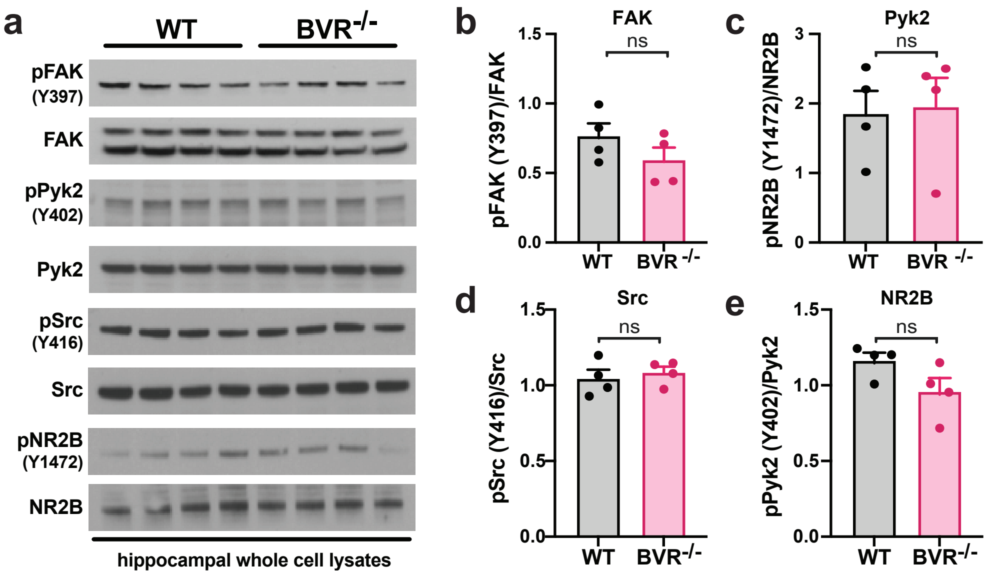


**Figure S2. Focal adhesion signaling is not globally disrupted in BVR-/- hippocampi.**

**a-e**, Immunoblots (*a*) and quantifications (*b*-*e*) of phospho-FAK (Y397), FAK, phospho-Pyk2 (Y402), Pyk2, phospho-Src (Y416), Src, phospho-NR2B (Y1472), and NR2B in hippocampal whole cell lysates from WT and $BVR^{-/-}$ mice. Data are expressed as normalized ratio of phosphorylated to total protein in each mouse. Points represent data from independent experiments. Mean ± SEM depicted. ns = $P > 0.05$, two-tailed unpaired Student's t-test.

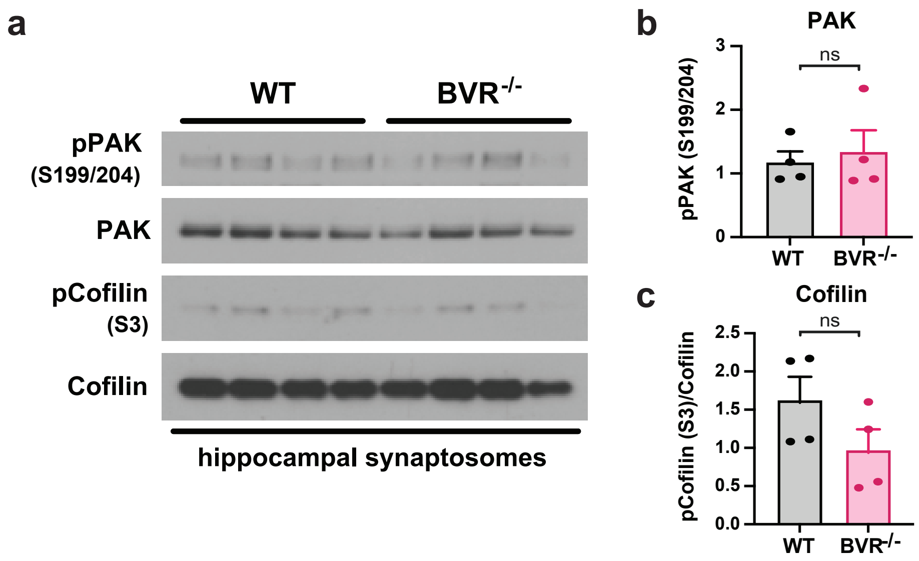


**Figure S3. Canonical FAK signaling is intact in BVR-/- synaptosomes.**

**a-c**, Immunoblots (*a*) and quantifications (*b*-*c*) of phospho-PAK (S199/204), PAK, phospho-Cofilin (S3), and Cofilin in crude synaptosomes isolated from hippocampi of WT and BVR$^{-/-}$ mice. Data are expressed as normalized ratio of phosphorylated to total protein in each mouse. Points represent data from independent experiments. Mean ± SEM depicted. ns = $P > 0.05$, two-tailed unpaired Student's t-test.

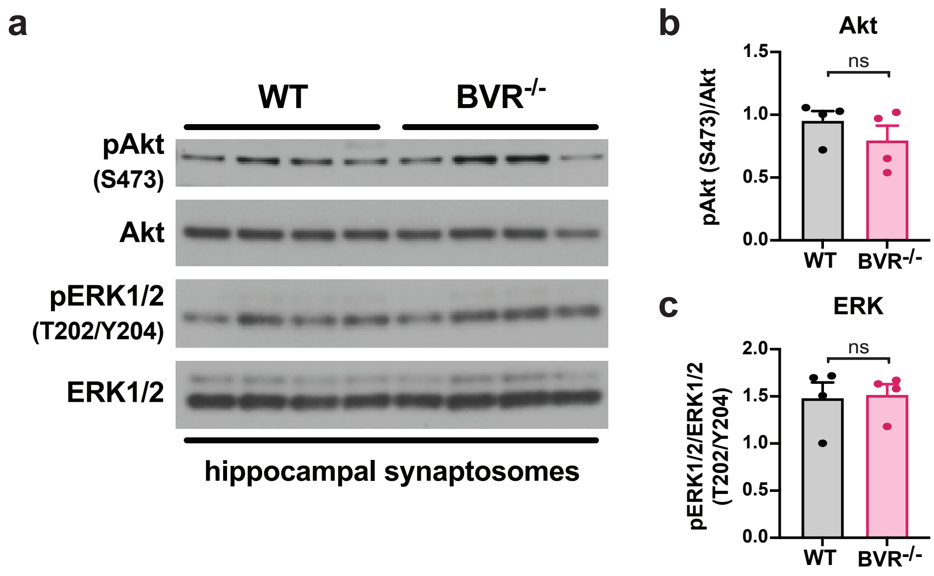


**Figure S4. PI3K/Akt and MAPK signaling is not disrupted in BVR$^{-/-}$ hippocampal synaptosomes.**

**a-c**, Immunoblots (*a*) and quantifications (*b-c*) of phospho-Akt (S473), Akt, phospho-ERK1/2 (T202/Y204), and ERK1/2 in crude synaptosomes isolated from hippocampi of WT and BVR$^{-/-}$ mice. Data are expressed as normalized ratio of phosphorylated to total protein in each mouse. Points represent data from independent experiments. Mean ± SEM depicted. ns = $P > 0.05$, two-tailed unpaired Student's t-test.

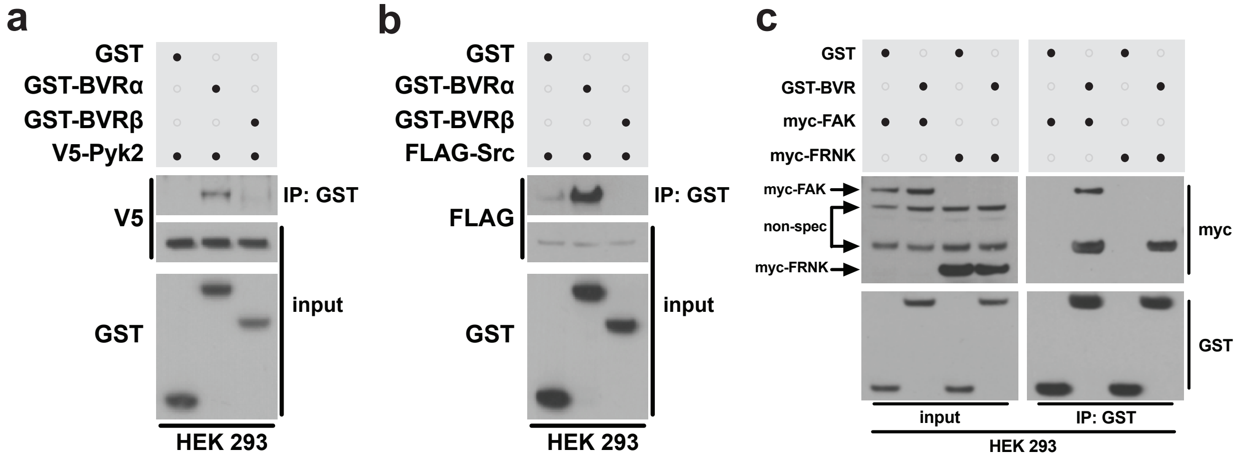


**Figure S5. BVR interaction with FAKs is isoform-specific.**

**a-b**, Immunoblots of lysates (input) and GST immunoprecipitates (IP) from HEK 293 cells overexpressing (*a*) V5-Pyk2 or (*b*) FLAG-Src and either GST, GST-BVRα, or GST-BVRβ. **c**, Immunoblots of lysates (input) and GST IP from HEK 293 cells overexpressing GST or GST-BVR and either myc-FAK or myc-FRNK. Immunoreactive bands corresponding to myc-FAK, myc-FRNK, or a nonspecific signal are indicated by labeled arrows.

**Table S1. Differential gene expression between WT and BVR-/- hippocampi.**

**Table S2. Genetic network analysis of neuroanatomic- and neural cell type- signatures downregulated in BVR-/- hippocampi.**

**Table S3. Gene ontology analysis of differential gene expression between WT and BVR-/- hippocampi.**
